# Inhibitory control of active expiration by the Bötzinger complex in rats

**DOI:** 10.1101/2019.12.19.883199

**Authors:** Karine C. Flor, William H. Barnett, Marlusa Karlen-Amarante, Yaroslav Molkov, Daniel B. Zoccal

## Abstract

The expiratory neurons of the Bötzinger complex (BötC) provide inhibitory inputs to the respiratory network, which, during eupnea, are critically important for respiratory phase transition and duration control. Herein, we investigated how the BötC neurons interact with the expiratory oscillator located in the parafacial respiratory group (pFRG) and control the abdominal activity during active expiration. Using the decerebrated, arterially perfused *in situ* rat preparations, we recorded the neuronal activity and performed pharmacological manipulations of the BötC and pFRG during hypercapnia or after the exposure to short-term sustained hypoxia – conditions that generate active expiration. The experimental data were integrated in a mathematical model to gain new insights in the inhibitory connectome within the respiratory central pattern generator. Our results reveal a complex inhibitory circuitry within the BötC that provides inhibitory inputs to the pFRG thus restraining abdominal activity under resting conditions and contributing to abdominal expiratory pattern formation during active expiration.

## INTRODUCTION

Oscillatory neural circuits are necessary components of the brain networks that sustain physiological and behavioral rhythms, such as sleep-wake cycle, hormone release, mastication, swallowing, locomotion and breathing (Shevtsova and Rybak, 2016, Saper, 2006, Kumar Jha et al., 2015, Harris-Warrick, 2010, Ramirez and Baertsch, 2018). In mammals, rhythmical contraction and relaxation of respiratory muscles emerges from interacting excitatory and inhibitory neurons with specific cellular properties, distributed within the pons and the medulla oblongata (Richter and Smith, 2014, Del Negro et al., 2018, Lindsey et al., 2012). Coupled oscillators embedded in this brainstem respiratory network are essential to generate and distribute synaptic inputs for the initiation of respiratory rhythmicity and the control of pattern formation (Anderson and Ramirez, 2017, Del Negro et al., 2018). Defining the arrangement and connections of the respiratory oscillators and circuitries are essential to understand how breathing is generated and adjusted to attend metabolic and behavior demands.

Overwhelming evidence suggests that a group of pre-inspiratory/inspiratory (pre-I/I) neurons with intrinsic bursting properties, located in the ventral surface of medulla, is a kernel of the circuit that generates inspiratory activity (Rybak et al., 2014, Rubin et al., 2011, Smith et al., 2007, Yang and Feldman, 2018, Kam et al., 2013, Phillips et al., 2019). These neurons are located in the so-called pre-Bötzinger complex (pre-BötC), and are able to endogenously generate rhythmic activity when isolated *in vitro* (Smith et al., 1991). Also, inhibition of pre-BötC *in vivo* causes persistent apnea (Tan et al., 2008), supporting the idea that the pre-BötC is a critical part of the respiratory oscillator. Rostral to the pre-BötC are found the expiratory neurons of the Bötzinger complex (BötC), which are important for the control of expiration (Ezure et al., 2003b, Tian et al., 1999, Smith et al., 2007). The BötC contains inhibitory (GABAergic and glycinergic) neurons with decrementing (post-inspiratory, post-I) or augmenting (aug-E) firing patterns during expiration (Bryant et al., 1993, Tian et al., 1999). They are suggested to establish mutual synaptic interactions with the pre-BötC neurons (Ezure et al., 2003c, Ezure et al., 2003a, Smith et al., 2007). Inhibition of BötC neurons, or disruption of inhibitory synapses in the BötC and/or pre-BötC dramatically depress breathing, suggesting that reciprocal inhibition between BötC and pre-BötC is necessary for respiratory rhythm regulation and generation of eupneic breathing (Marchenko et al., 2016, Bongianni et al., 2010, Fortuna et al., 2018, Ezure, 1990). The BötC and pre-BötC neurons, therefore, are proposed to constitute the core of the respiratory network (Richter and Smith, 2014).

Under resting conditions, the mammalian breathing exhibits a three-phase pattern that includes inspiration (contractions of inspiratory pumping muscles), post-inspiration (first stage of expiration where expiratory flow occurs passively but is regulated by upper airway muscles) and passive expiration (second stage of expiration) (Del Negro et al., 2018, Richter and Smith, 2014, Bianchi et al., 1995). In states of elevated metabolic demand, such as during physical exercise or blood gas challenges (e.g. hypoxia and hypercapnia), contractions of abdominal expiratory muscles develop mostly during the second stage of expiration to forcefully exhale the air from the lungs, hence increasing pulmonary ventilation (Jenkin and Milsom, 2014, Lemes and Zoccal, 2014). This pattern of active expiration emerges due to the recruitment of the conditional expiratory oscillator located rostral to the BötC and ventrolateral to the facial nucleus, within the parafacial respiratory group (pFRG) (Janczewski and Feldman, 2006, Abdala et al., 2009). The pFRG contains expiratory neurons that are silent under normoxia and normocapnia, but fire rhythmically during the late phase of expiration (late-E) in conditions of hypoxia or hypercapnia (Abdala et al., 2009, Moraes et al., 2012a, Molkov et al., 2010). Pharmacological or optogenetic suppression of the pFRG neurons eliminates the abdominal expiratory activity evoked by hypercapnia or stimulation of peripheral chemoreceptors (Zoccal et al., 2018, de Britto and Moraes, 2017, Marina et al., 2010, Moraes et al., 2012a, Lemes et al., 2016), indicating that pFRG neurons are necessary for the emergence of active expiratory pattern. How pFRG oscillator interacts with other respiratory compartments within the brainstem, especially with the core of respiratory network, was theoretically hypothesized (Molkov et al., 2016, Molkov et al., 2014, Rubin et al., 2011, Molkov et al., 2010), but largely remains an unresolved question.

Although the pFRG is partially overlapped with the chemosensitive retrotrapezoid nucleus (RTN) (Guyenet, 2014), recent evidence indicates that expiratory neurons of the pFRG are not sensitive to CO_2_/pH (de Britto and Moraes, 2017) and require excitatory inputs to be activated (Zoccal et al., 2018). Part of the excitatory drive to the pFRG arises in the RTN and the pre-BötC neurons (Zoccal et al., 2018, Huckstepp et al., 2016, Barnett et al., 2017). Inhibitory synapses are also needed for the proper control of the pFRG activity, hence the pattern formation of abdominal motor activity (Pagliardini et al., 2011, Barnett et al., 2018). Under resting conditions, the pFRG oscillator is synaptically suppressed, and its pharmacological disinhibition (by the application of antagonists for GABAergic and glycinergic receptors) generates active expiration under normoxia and normocapnia (Pagliardini et al., 2011, de Britto and Moraes, 2017). *In silico* studies propose that the inhibitory drive to the pFRG originates, at least in part, from the ventral respiratory column (Molkov et al., 2010). Specifically, computational modeling suggests that neurons of the BötC might be a source of inhibition to the pFRG oscillator, and the reduction of this inhibitory drive might be an important stage for the emergence of active expiration in conditions of elevated excitatory drive (Barnett et al., 2018, Jenkin et al., 2017). However, experimental and functional evidence is still required to confirm the possible inhibitory control of the BötC neurons over the pFRG expiratory oscillator.

In the present study, we investigated the role of the BötC neurons in the generation of active expiratory pattern in rats. Firstly, we characterized the dynamic changes that occur in the activity of the BötC neurons (aug-E and post-I) during the emergence of active expiration. Using pharmacological approaches to promote stimulation or disinhibition, we then tested the hypothesis that the BötC neurons provides inhibitory inputs to the pFRG that critically control the generation of active expiration in conditions of elevated metabolic demand. The experimental data were combined with an extended mathematical modeling to explore and functionally describe an inhibitory circuitry within the BötC in two experimental models of active expiration: i) acute exposure to hypercapnia (Abdala et al., 2009, Molkov et al., 2011) and ii) after the exposure to short-term (24 h) sustained hypoxia, which evokes sustained active expiration under resting conditions (Flor et al., 2018, Moraes et al., 2014). In both conditions, the pFRG expiratory oscillator is recruited and abdominal motor hyperactivity is observed (Abdala et al., 2009, Moraes et al., 2014).

## RESULTS

### Discharge pattern of post-I and aug-E neurons of BötC during normocapnia and hypercapnia in control in situ preparations

Under normocapnic conditions, *in situ* preparations of control rats exhibited an eupneic-like respiratory pattern, with ramping PN bursts, low AbN activity during E1 and E2 phases, and inspiratory/post-inspiratory components in cVN (Figures 1 and 2). In the ventrolateral medulla of these preparations, within the BötC region, we identified two types of expiratory neurons: i) neurons that exhibited a peak of discharge during early E1 phase, immediately after PN burst, followed by a decrementing pattern of discharge during the expiratory phase, named post-I neurons (n=5 cells from 4 *in situ* preparations, Figure 1A and B); and ii) neurons with an augmenting pattern of discharge, with maximal firing frequency during E2 phase, named aug-E neurons (n=9 cells from 7 *in situ* preparations, Figure 2A and B). These neurons were found at the level of −12.12 and −12.48 mm relative to Bregma (Paxinos and Watson, 2007), 1726±80 mm lateral to midline and 433±58 from ventromedullary surface.

**Figure 1.**
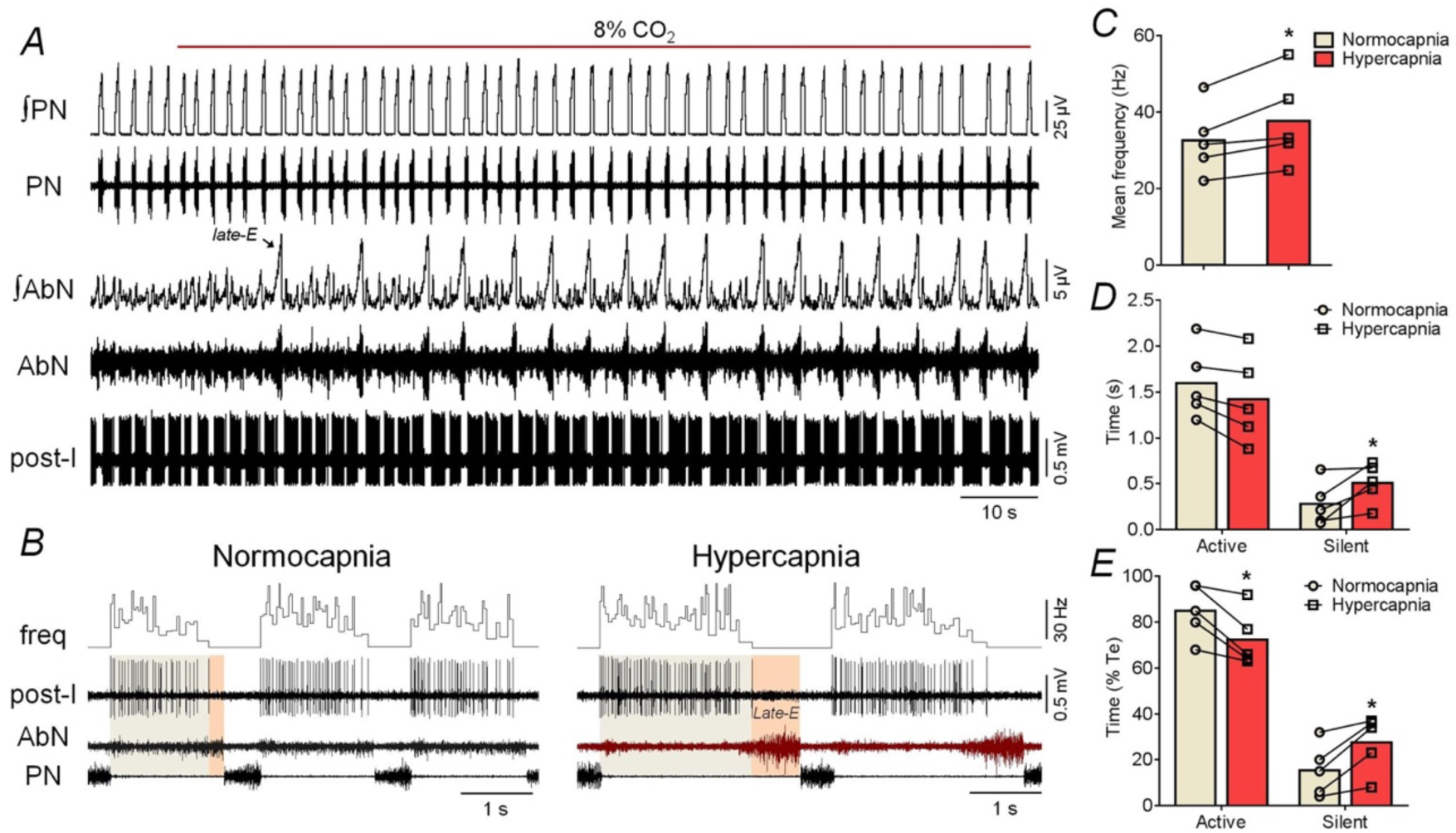
Changes in the discharge pattern of post-I neurons in the BötC during the exposure to hypercapnia. **Panel A:** raw and integrated (∫) recordings of phrenic (PN) and abdominal (AbN) nerve activities, and unitary recordings a post-I neuron of the BötC of a control *in situ* preparation, representative from the group, illustrating the emergence of active expiration (late-E bursts in AbN, arrows) with the increase of fractional concentration of CO_2_ to 8% in the perfusate. **Panel B**: expanding recordings from panel A, illustrating the pattern of PN, AbN and post-I neuronal activities (unit and instantaneous frequency) under normocapnia and hypercapnia. The colored boxes delineate the active and silent periods of neuronal activity. Note the absence of action potentials in post-I recordings when AbN late-E burst emerges. **Panels C-E**: average values of mean firing frequency (C), and durations of active and silent periods of BötC post-I neurons, in seconds (D) and in percentage values relative to the expiratory time (E), during normocapnia and hypercapnia (n=5). * different from corresponding values during normocapnia, P<0.05.

**Figure 2.**
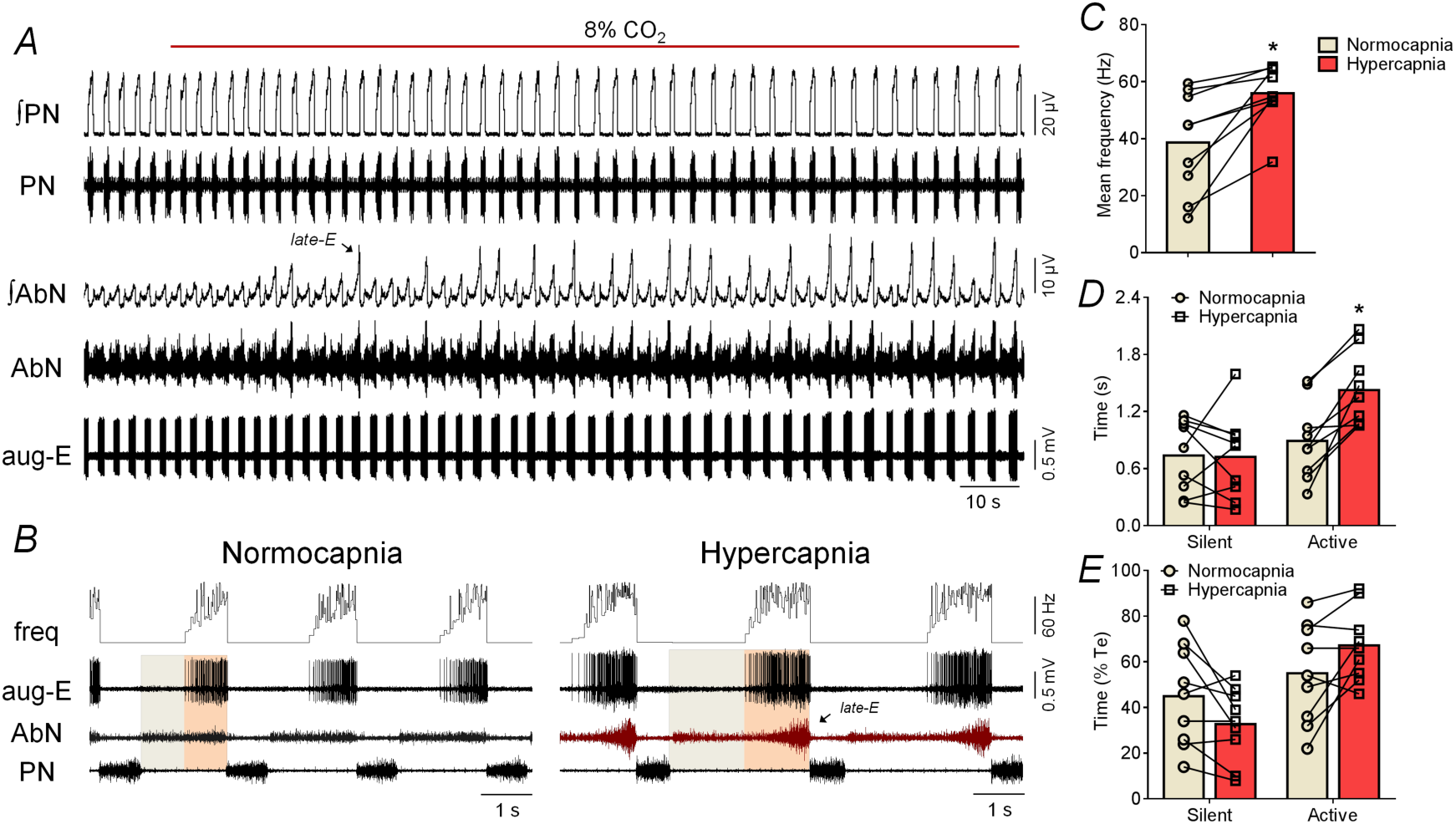
Changes in the discharge pattern of aug-E neurons in the BötC during the exposure to hypercapnia. **Panel A:** raw and integrated (∫) recordings of phrenic (PN) and abdominal (AbN) nerve activities, and unitary recordings an aug-E neuron of the BötC of a control *in situ* preparation, representative from the group, illustrating the emergence of active expiration (late-E bursts in AbN, arrows) with the increase of fractional concentration of CO_2_ to 8% in the perfusate. **Panel B**: expanding recordings from panel A, illustrating the pattern of PN, AbN and aug-E neuronal activities (unit and instantaneous frequency) under normocapnia and hypercapnia. The colored boxes delineate the active and silent periods of neuronal activity. Note the increased firing frequency of aug-E neuron when AbN late-E burst emerges. **Panels C-E**: average values of mean firing frequency (C), and durations of active and silent periods of BötC aug-E neurons, in seconds (D) and in percentage values relative to the expiratory time (E), during normocapnia and hypercapnia (n=9). * different from corresponding values during normocapnia, P<0.05.

The increase in the fractional concentration of CO_2_ in the perfusate modified the respiratory pattern and generated active expiration, as well as changed the activity of BötC neurons in control *in situ* preparations (Figures 1, 2 and 3). With respect to motor outputs (n=7), we observed that hypercapnia: i) decreased PN burst frequency, ii) did not modify PN burst amplitude, iii) reduced Ti and prolonged Te, iv) did not change cVN post-inspiratory duration and mean activity, and v) did not modify AbN activity during E1 phase, but markedly increased during E2 phase due to the emergence of bursts during the late part of expiration (late-E). These responses are summarized on Table 3. In association with the respiratory motor changes, hypercapnia promoted a modest but significant increase in the average firing frequency of post-I during E1 phase (32.6±9.1 vs 37.8±11.8 Hz, P=0.0253; Figures 1A-C). Moreover, the pattern of post-I neuronal discharge during hypercapnia showed a negative relationship with the emergence of active expiration, with either absence or very low firing frequency during the occurrence of AbN late-E bursts in the E2 phase (Figure 1B). As a result, the duration of active time of post-I neurons during hypercapnia significantly reduced (1.599±0.392 vs 1.422±0.478 s, P=0.0615, Figure 1D; 85±12 vs 72±12 % of expiratory time, P=0.0097, Figure 1E) while the silent period prolonged (0.280±0.240 vs 0.510±0.219 s, P=0.0185, Figure 1D; 15±11 vs 28±12 % of expiratory time, P=0.0116, Figure 1E). The exposure to hypercapnia also increased the firing frequency of aug-E neurons during E2 phase (44.1±18.4 vs 54.6±10.0 Hz, P=0.0039; Figures 2A-C). On the other hand, different from post-I population, the aug-E neurons displayed a positive association with hypercapnia-induced active expiration (Figure 2B), with a prolonged active time during E2 phase (0.891±0.409 vs 1.425±0.391 s, P=0.0178, Figure 2D; 55±22 vs 68±16 % of expiratory time, P=0.0727, Figure 2E), with no significant changes in the duration of silent period during E1 phase (0.739±0.377 vs 0.724±0.447 s, P>0.99, Figure 2D; 45±22 vs 33±16 % of expiratory time, P=0.0727, Figure 2E). These findings indicate that although BötC post-I and aug-E exhibited an increased firing frequency under hypercapnia, these neurons are distinctly controlled during the occurrence of abdominal bursts in the late-E phase.

**Figure 3.**
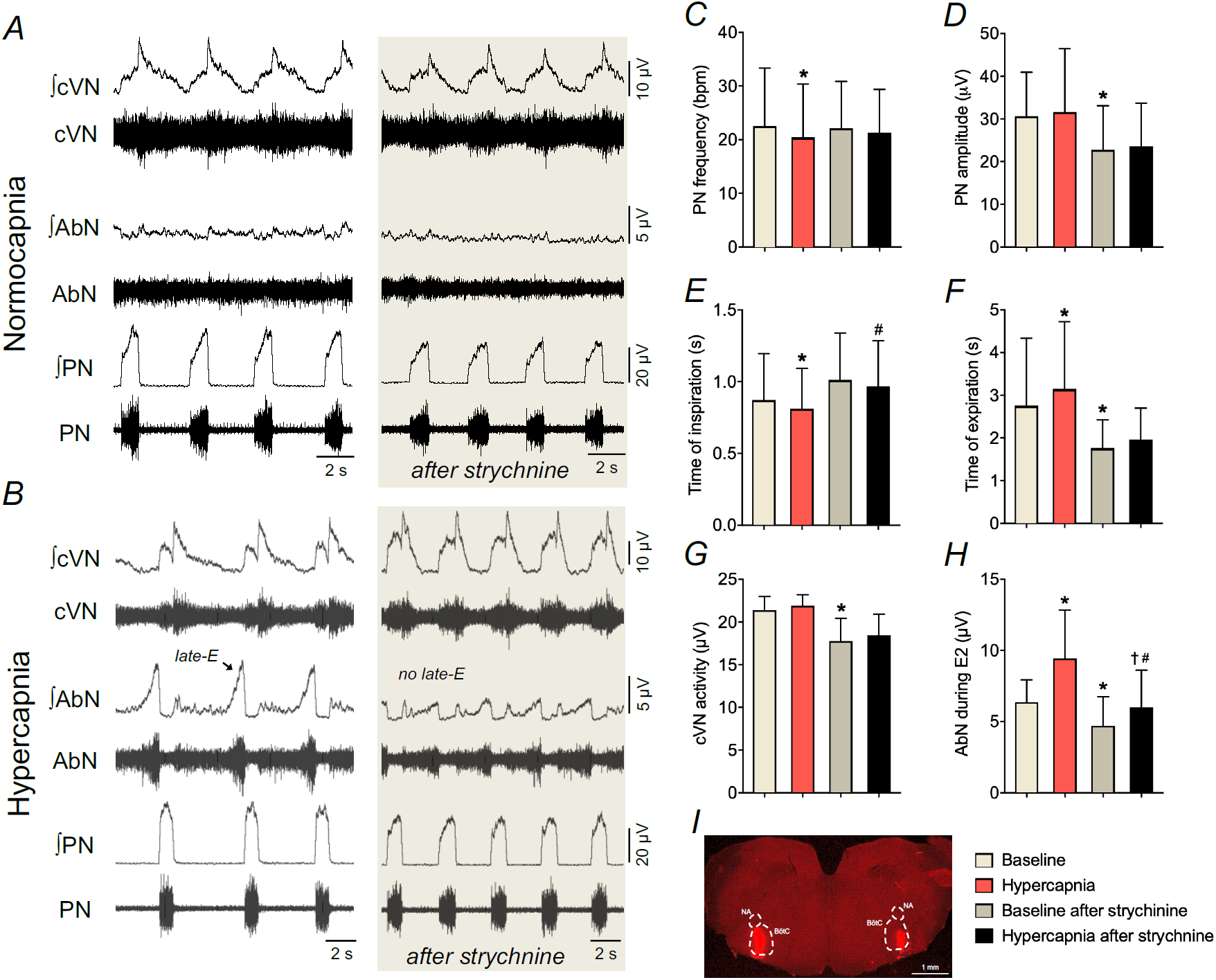
Strychnine microinjections in the BötC suppress the emergence of active expiration during hypercapnia. **Panels A-B:** raw and integrated (∫) recordings of cervical vagus (cVN), abdominal (AbN) and phrenic (PN) nerve activities of a control *in situ* preparation, representative from the group, under normocapnia and hypercapnia, respectively, before and after bilateral microinjections of strychnine (10 µM) in the BötC. **Panels C-H**: average values of PN frequency and amplitude, times of inspiration and expiration, cVN post-I and AbN E2 activities, respectively, during baseline (normocapnia) and hypercapnic conditions, before and after strychnine microinjections in the BötC of control *in situ* preparations (n=7). * different from baseline, # different from baseline after strychnine, † different from hypercapnia, P<0.05. **Panel I**: coronal section of brainstem from a control *in situ* preparation, illustrating the sites of bilateral microinjections of strychnine in the BötC.

### Distinct effects of glycinergic and GABAergic transmission within the Bötzinger complex on the control of abdominal activity during normocapnia and hypercapnia

To verify the role of inhibitory synapses within the BötC on the formation of AbN activity in control animals, we initially explored the effects of glycine receptor antagonism with microinjections of strychnine (10 µM) in the BötC of control *in situ* preparations (n=7). The concentration of strychnine used in our study was smaller than described previously that caused disruption of respiratory pattern when microinjected in the BötC/pre-BötC of *in situ* preparations (Marchenko et al., 2016). Under normocapnic conditions, strychnine microinjections in the BötC slightly reduced the amplitude of PN, cVN and AbN outputs as well as the durations of expiration and the post-I component of cVN. No significant changes were observed in the PN burst frequency and time of inspiration (Figures 3A, 3C-H, Table 3). During hypercapnia, strychnine microinjections in the BötC was able to prevent the changes in the PN frequency (Figure 3C) and time of expiration (Figure 3F), but not the alterations in the PN amplitude (Figure 3D), time of inspiration (Figure 3E), cVN post-inspiratory activity (Figure 3G) and duration (Table 3). Moreover, disinhibition of BötC neurons markedly attenuated the hypercapnia-induced late-E bursts in AbN (Figure 3H). Bilateral microinjections of vehicle in the BötC of *in situ* preparations of control rats (n=3) did not modify the respiratory motor activities either under baseline conditions or in response to hypercapnia (data not shown). The sites of microinjections in the BötC are illustrated in Figure 3I. These findings indicate that glycinergic transmission within the BötC controls neurons whose activation is sufficient to inhibit the emergence of active expiration during hypercapnia.

In another set of experiments (n=7), we examined the effects of reduction of GABAergic neurotransmission within the BötC, with bilateral microinjections of gabazine (250 µM) (Barnett et al., 2018), on the abdominal activity of control *in situ* preparations (Figure 4). Different from strychnine microinjections, gabazine in the BötC elicited the pattern of active expiration under normocapnic conditions (Figure 4A and B). Specifically, the antagonism of GABAergic synapses in the BötC caused: i) reductions in the PN burst frequency (29±5 vs 22±5 cpm, P=0.0028, Figure 4C), but not in the amplitude (26±7 vs 28±8 µV, P=0.0973; Figure 4D); ii) no changes in the inspiratory duration (0.708±0.141 vs 0.791±0.241 s, P=0.1931; Figure 4E), but a significant increase in the expiratory duration (1.403±0.313 vs 2.096 ± 0.412 s, P=0.0027, Figure 4F); iii) no changes in the cVN post-inspiratory mean activity (15.9±1.8 vs 16.9±1.9 µV, P=0.2775; Figure 4G) and normalized duration (70±17 vs 67±15 % of Te, P=0.3505); and vi) no significant changes in the AbN activity during E1 phase (1.9±0.7 vs 2.4±1.2 µV, P=0.1684), but a markedly increase in the AbN activity during E2 phase (3.2±2.1 vs 6.5±3.8 µV, P=0.0061, Figure 4H) due to the emergence of *late-E* bursts (Figure 4B). The exposure to hypercapnia after gabazine microinjections in the BötC did not cause any additional effects on nerve outputs (data not shown). The sites of microinjections in the BötC are represented in Figure 4I. These findings indicate that GABAergic transmission within the BötC affects neurons whose activation can inhibit neurons restraining active expiration during normocapnia.

**Figure 4.**
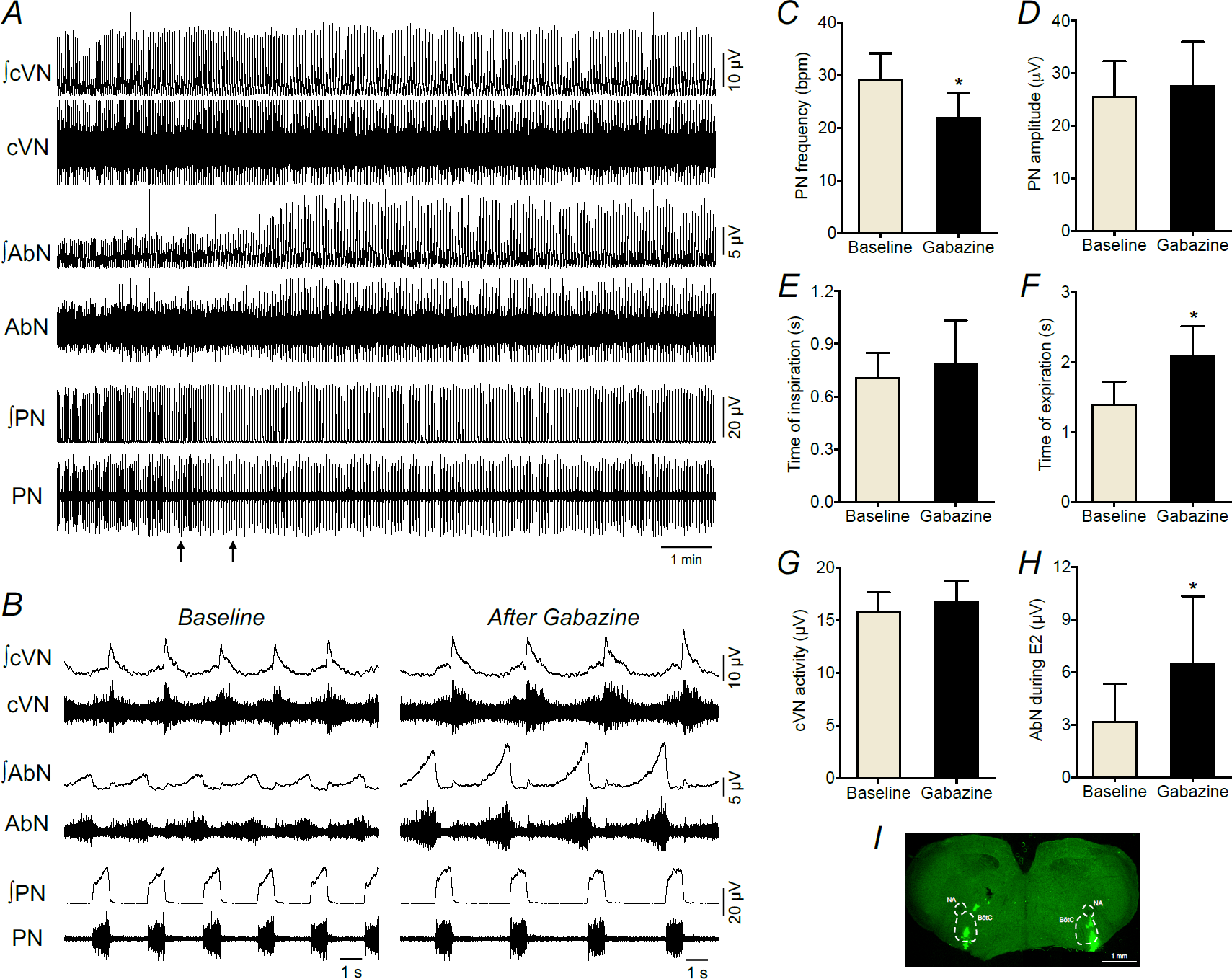
Suppression of GABAergic transmission in the BötC evokes active expiration under normocapnic conditions. **Panel A:** raw and integrated (∫) recordings of cervical vagus (cVN), abdominal (AbN) and phrenic (PN) nerve activities of a control *in situ* preparation under normocapnia, representative from the group, before and after bilateral microinjections of gabazine (250 µM) in the BötC (arrows). **Panel B:** Expanded tracings from panel A, demonstrating the respiratory pattern before (baseline) and after gabazine microinjections. **Panels C-H**: average values of PN frequency and amplitude, times of inspiration and expiration, cVN post-I and AbN E2 activities, respectively, before (baseline) and after gabazine microinjections in the BötC of control *in situ* preparations (n=7). * different from baseline, P<0.05. **Panel I**: coronal section of brainstem from a control *in situ* preparation, illustrating the sites of bilateral microinjections of gabazine in the BötC.

### Proposed model of GABAergic and glycinergic circuitry within the Bötzinger complex that supports differential respiratory responses to gabazine and strychnine

To explain the opposite effects of GABAergic and glycinergic neurotransmission in the BötC on the emergence of active expiration, we propose a local GABAergic/glycinergic circuitry within the BötC. Specifically, we assumed that the post-inspiratory populations are formed by GABAergic and glycinergic cells. These neuronal groups established mutual interactions with the pFRG, where the post-I_GABA_ and post-I_Gly_ populations sent projections to the late-E population of the pFRG, whilst the BötC aug-E neurons received excitatory inputs from the pFRG population (Figure 5). In normocapnia, late-E neurons of the pFRG were not active, which was accomplished in the model by phase-spanning inhibition due to inhibition from populations in the BötC as well as the early-I population of the pre-BötC (Figure 5). The pacemaker-like pre-I/I and the adapting early-I populations composed the inspiratory neuronal populations of the pre-BötC and were responsible for the initiation of the inspiratory phase (Figure 6A). At the transition from inspiration to expiration, the post-I_GABA_ and post-I_Gly_ neurons of the BötC fired strongly at first and then waned. The model was constructed such that the neurons of the post-I_GABA_ population possessed a slow outward current that provided an adaptation mechanism, where the firing rate reduced steadily during the expiratory phase in the absence of inhibitory input. On the other hand, the activity of the post-I_Gly_ population was shaped by inhibition from aug-E population: as aug-E incremented over the course of expiration, post-I_Gly_ was significantly suppressed (Figure 6A).

**Figure 5.**
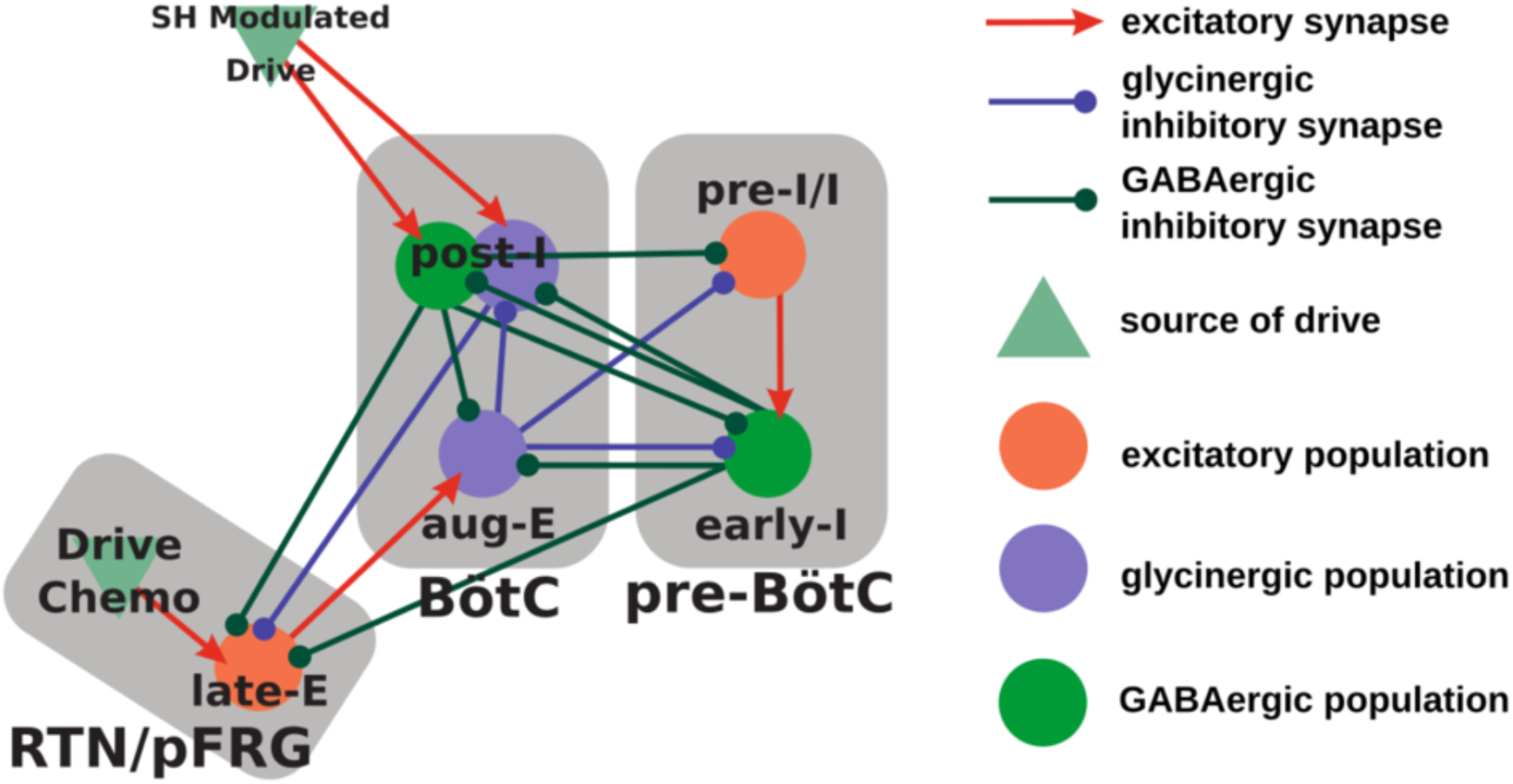
Model connectivity schematic includes neuronal populations in the pre-Bötzinger (pre-BötC) and the Bötzinger complexes (BötC) as well as the retrotrapezoid nucleus (RTN) and the parafacial respiratory group (pFRG). This respiratory circuitry includes the pre-inspiratory/inspiratory (pre-I/I) population and the early-inspiratory population (early-I) of the pre-BötC; the augmenting-expiratory population (aug-E) and the post-inspiratory population (post-I) of the BötC, and the late-expiratory population (late-E) of the pFRG.

**Figure 6.**
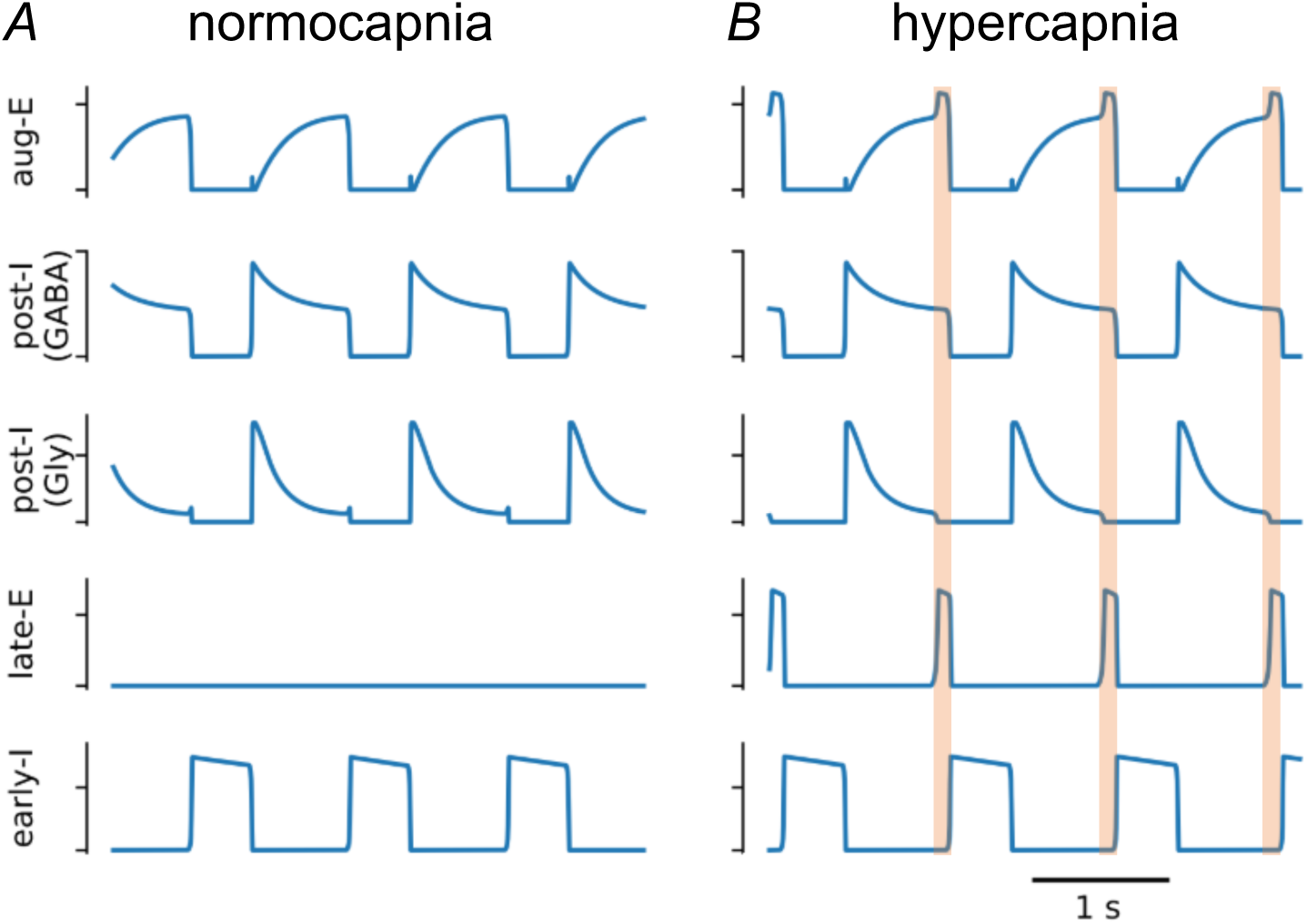
Model simulations of neuronal activity during normocapnia and hypercapnia. Active expiration emerges in model activity following an increase in chemosensitive drive to the late expiratory population (late-E) of the parafacial respiratory group (pFRG). The baseline model activity (**Panel A**) transforms to include late-expiratory bursts (**Panel B**) when the chemosensitive drive to late-E is increased by 33%. The transformation of the respiratory pattern during the late expiratory phase is emphasized by the orange shaded boxes. Other abbreviations – aug-E: augmenting expiratory population; post-I (GABA): GABAergic post-inspiratory population; post-I (Gly): glycinergic post-inspiratory population; early-I: early inspiratory population.

Hypercapnia was implemented as an increase of 33% in the chemosensitive drive to late-E neurons of the pFRG. The emergence of active expiration occurred when this drive was sufficient for the late-E population to overcome expiratory inhibition from the GABAergic and glycinergic post-I populations (Figure 6B). Since this expiratory inhibition was decrementing, late-E bursts occurred at the end of expiration when inhibition was weakest. Moreover, late-E activity facilitated aug-E activity, which then terminated post-I activity during the late-E burst (Figure 6B). During inspiration, early-I inhibitory drive was able to suppress late-E activity in the pFRG.

Based on these model conditions, we then simulated the experiments with gabazine microinjections in the BötC under normocapnia by reducing the synaptic weights from the early-I to post-I_Gly_, early-I to post-I_GABA_, and early-I to aug-E populations by 10% each. Besides, the post-I_GABA_ to aug-E synaptic weight was reduced by 15%. The chemosensitive drive was not changed. Following these manipulations, the model exhibited late-E bursts at the end of expiration similar to the activity observed in experimental recordings (Figure 7A and B). Although a number of synaptic weights were altered to reproduce this experimental condition, the change in the weight of the post-I_GABA_ to aug-E synapse was the main factor responsible for respiratory pattern transformation. Early-I is only active during inspiration, so the change to its weights did not directly alter inhibitory tone in the pFRG during expiration. The decrease in the weight of synaptic inhibition from post-I_GABA_ to aug-E resulted in a facilitation of aug-E firing rate. Aug-E inhibition of post-I_Gly_ was greater at the end of expiration, and the post-I_Gly_ firing rate was sufficiently suppressed to release late-E population of the pFRG from glycinergic inhibition (Figure 7B). Late-E activation transiently facilitated aug-E, which is the mechanism that supports the gap between the termination of post-I activity and the expiration-to-inspiration phase transition.

**Figure 7.**
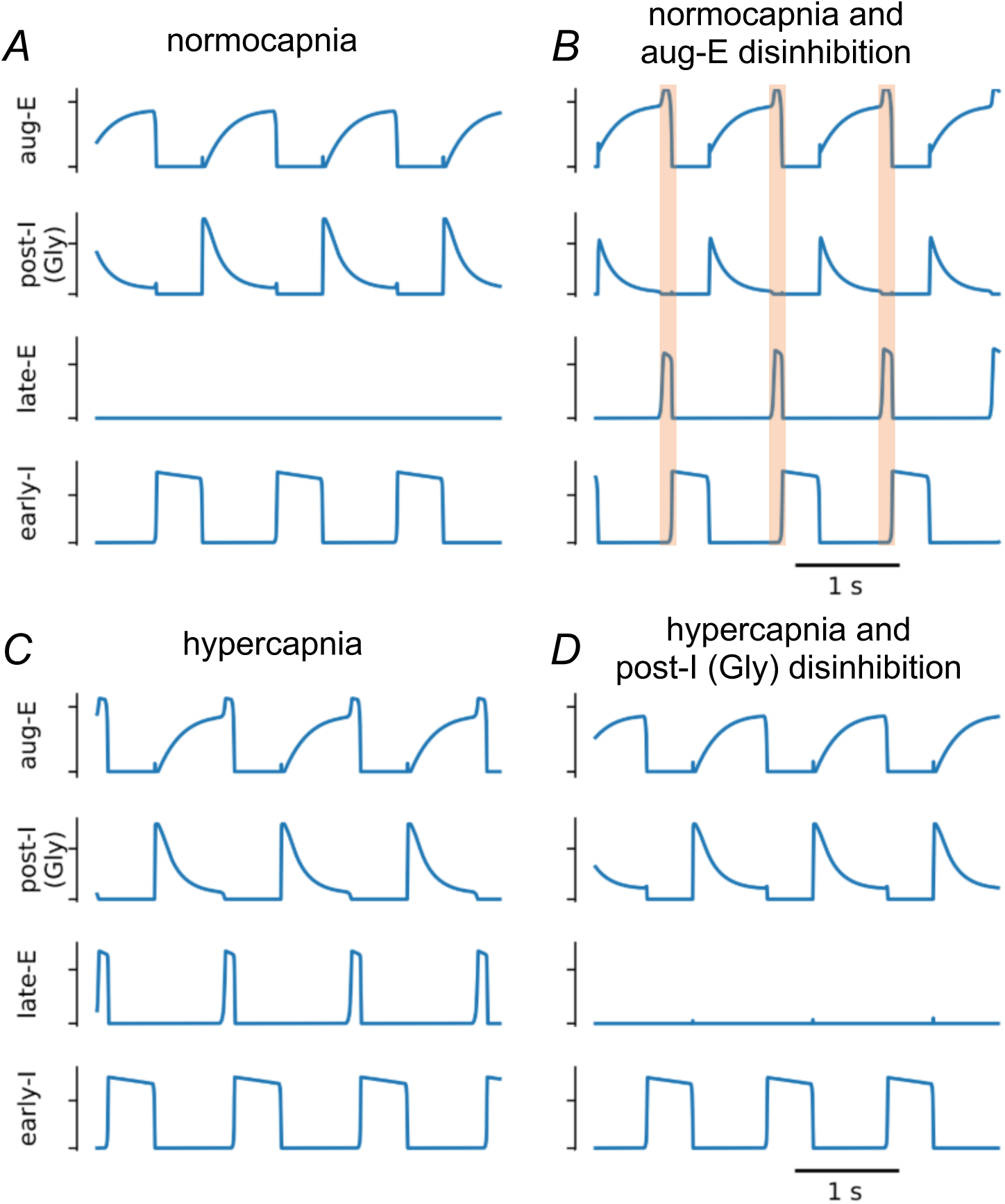
Model simulations of GABAergic and glycinergic disruptions in the Bötzinger complex. **Panels A and B:** simulations of reduced GABAergic transmission within the BötC generated active expiration under baseline conditions. The baseline model activity (**Panel A**) is transformed to exhibit late expiratory activity (**Panel B**) following the reduction of the early-inspiratory population (early-I) to the glycinergic post-inspiratory population [post-I (Gly)] synaptic weight, the early-I to GABAergic post-inspiratory population [post-I (GABA)] synaptic weight, and the early-I to augmenting expiratory (aug-E) synaptic weight by 10% as well as the post-I (GABA) to aug-E synaptic weight by 15%. The main factor contributing to the expiratory pattern transformation was the change in the post-I (GABA) to aug-E synaptic weight. The transformation of the respiratory pattern during the late expiratory phase is emphasized by the orange shaded boxes. **Panels C and D**: In simulations with increased chemosensitive drive to the late expiratory population (late-E), the reduction of glycinergic transmission within the BötC suppressed the active expiration. Active expiration was induced by increasing the chemosensitive drive to late-E by 33% (**Panel C**), and then in a follow-up simulation active expiration was abolished by reducing the weight of the inhibitory synapse from the aug-E population to the post-I (Gly) population (**Panel D**).

To simulate the scenario of strychnine microinjections in the BötC during hypercapnia, we reduced synaptic weights of the glycinergic circuitry within the BötC (Figure 7 C and D). The weight of the aug-E to post-I_Gly_ synapse was reduced by 10%. This manipulation was sufficient to suppress the activity of the late-E population of the pFRG recruited by the increase in the chemosensitive drive (hypercapnia condition). The mechanism responsible for this transformation of the respiratory pattern revolves around the disinhibition of the glycinergic post-I population. Our manipulation reduced the inhibitory output of aug-E, which facilitated the firing rate of post-I_Gly_ neurons. This change in firing rate directly resulted in greater glycinergic inhibition on late-E neurons of the pFRG (Figure 7D).

### Active expiration after the exposure to sustained hypoxia in situ and in silico

In order to broaden our understanding about active expiration in different conditions, and to gain insights into Bötzinger complex and parafacial respiratory group interactions, we also performed experiments using the experimental model of rats exposed to short-term sustained hypoxia (10% O_2_, 24 h). In agreement with previous studies (Flor et al., 2018, Moraes et al., 2014), rats previously exposed to (n=9) exhibited the pattern of active expiration under conditions of normocapnia (Figure 8A). In comparison to control group (n=9), the *in situ* preparations of SH rats displayed: i) reduced PN burst frequency (28±9 vs 19± 4 bpm, P=0.0160, Figure 8D); ii) no changes in the PN burst amplitude (29.7±12.1 vs 29.5±11.1 µV, P=0.9654) and time of inspiration (0.689±0.167 vs 0.714±0.192 s, P=0.7699, Figure 8E); iii) prolonged time of expiration (1.712±0.765 vs 2.629±0.761 s, P=0.0214, Figure 8F); iv) augmented cVN post-inspiratory mean activity (11.3±8.4 vs 20.2±8.8 µV, P=0.0445, Figure 8G); and v) AbN hyperactivity during E2 phase (2.3±1.3 vs 6.2±3.4 µV, P=0.0092, Figure 8H) due to the presence of late-E bursts (Figure 8A). Consistent with previous observations (Moraes et al., 2014), we also verified that the BötC post-I neurons of SH rats (n=2 cells from 2 preparations) stopped firing at the onset of AbN late-E bursts (Figure 8B), indicating a negative relationship between post-I neuronal activity and AbN late-E breakthrough in this experimental model. On the other hand, the BötC aug-E neurons of SH (n=2 cells from 2 preparations) exhibited prolonged activity in the presence of AbN late-E activity (Figure 8C), suggesting a positive relationship after SH exposure. The average frequency of discharge of post-I and aug-E neurons of SH rats *in situ* were 49.7±15.4 and 63.0±2.8 Hz, respectively. This pattern of activity of post-I and aug-E neurons of SH rats resembled the activity of BötC neurons observed in control rats during hypercapnia (Figures 1 and 2).

**Figure 8.**
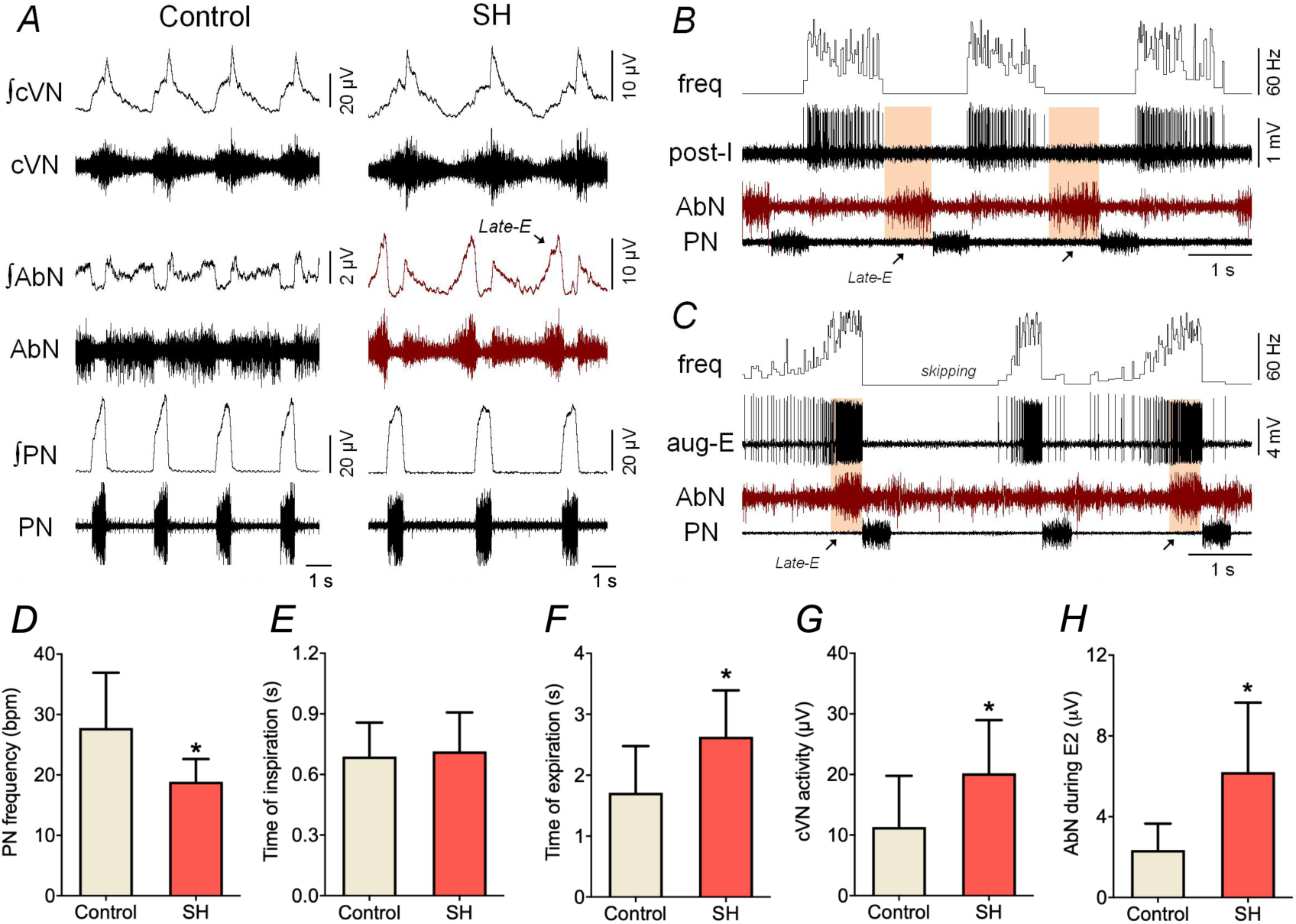
Respiratory motor pattern and BötC neuronal activity in rats exposed to sustained hypoxia. **Panel A:** raw and integrated (∫) recordings of cervical vagus (cVN), abdominal (AbN) and phrenic (PN) nerve activities of control and SH *in situ* preparations, representative from their groups, under normocapnia. Note that SH preparations exhibit AbN late-E bursts (active expiration) at resting conditions. **Panels B-C**: recordings of post-I and aug-E neurons (unit and instantaneous frequency), respectively, a two *in situ* preparations of SH rats, representative from the group, illustrating the relationship between BötC neuronal activity and active expiration. In panel B, note the absence of action potentials in post-I activity when late-E burst emerged. In panel C, note the increased firing frequency of aug-E neurons in the presence of AbN late-E burst, in comparison to a respiratory cycle when AbN late-E burst skipped (skipping). **Panels D-H**: Average values of PN frequency, times of inspiration and expiration, and cVN post-I and AbN E2 activities of *in situ* preparations of control (n=9) and SH rats (n=9). * different from control group, P<0.05.

Considering the discharge profile of post-I and aug-E neurons in SH rats, we then simulated the SH condition by reducing the excitability of neurons that fire most strongly following the inspiration-to-expiration phase transition (Figure 9 A and B). In our model, both GABAergic and glycinergic post-I populations of the BötC inhibit the late-E population of the pFRG, and a decrease in the conductance of their tonic excitatory drive by 5% accomplished a sufficient decrease in their firing rates at the end of expiration and, hence, disinhibition of the late-E population to a sufficient extent to induce active expiration under normal chemosensitive drive (Figure 9 B). Based upon our simulations that the SH-induced active expiration is closely associated with the reduction of excitability of post-inspiratory neurons, one can anticipate that the abdominal hyperactivity after SH exposure can be abolished by a uniform increase in the excitability of neurons in the BötC. Using the SH model, we tested this scenario by introducing an additional tonic excitatory conductance, and it was implemented as a transient 12% increase in the total conductance of excitatory drive in each of the three neuronal populations in the BötC. This manipulation eliminated the activity of late-E neurons, as demonstrated in the Figure 10A. In simulation, the firing rate of the post-I_GABA_, aug-E and post-I_Gly_ populations increased during the epoch with enhanced excitation (Figure 10A). Since post-I_GABA_ and post-I_Gly_ both project to late-E, we can formulate a testable prediction that a uniform increase in excitatory conductance in the BötC would result in greater expiratory inhibition on the late-E population of the pFRG that would prevent the latter from activation normally observed after SH exposure.

**Figure 9.**
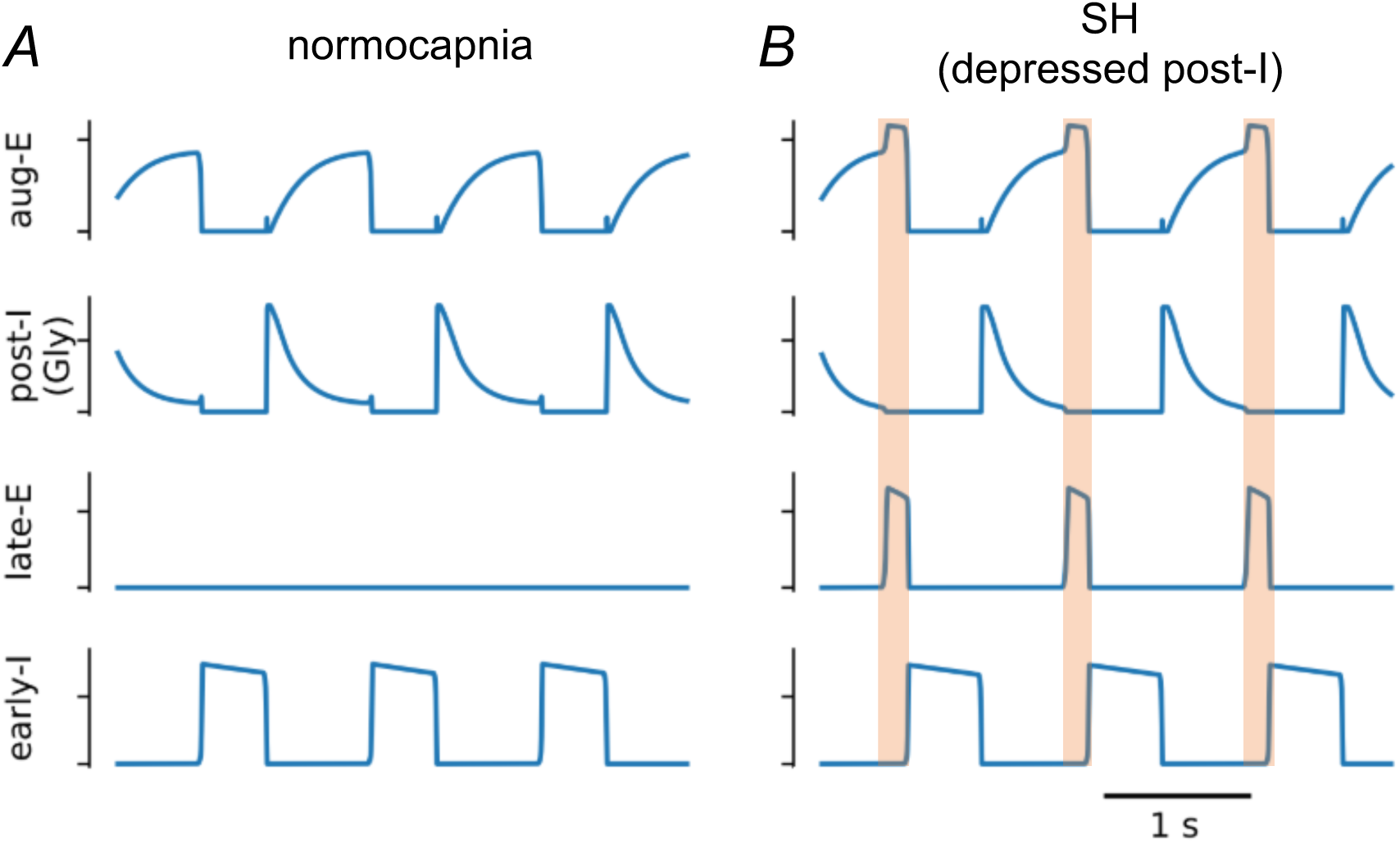
Model simulations of active expiration induced by sustained hypoxia. The baseline three phase rhythm (**Panel A**) was transformed to pattern that included active expiration (**Panel B**) by reducing excitatory conductances on the GABAergic and glycinergic post-inspiratory (post-I) neuronal populations by 5%, which caused the recruitment of late-expiratory (late-E) population. The transformation of the respiratory pattern during the late expiratory phase is emphasized by the orange shaded boxes.

**Figure 10.**
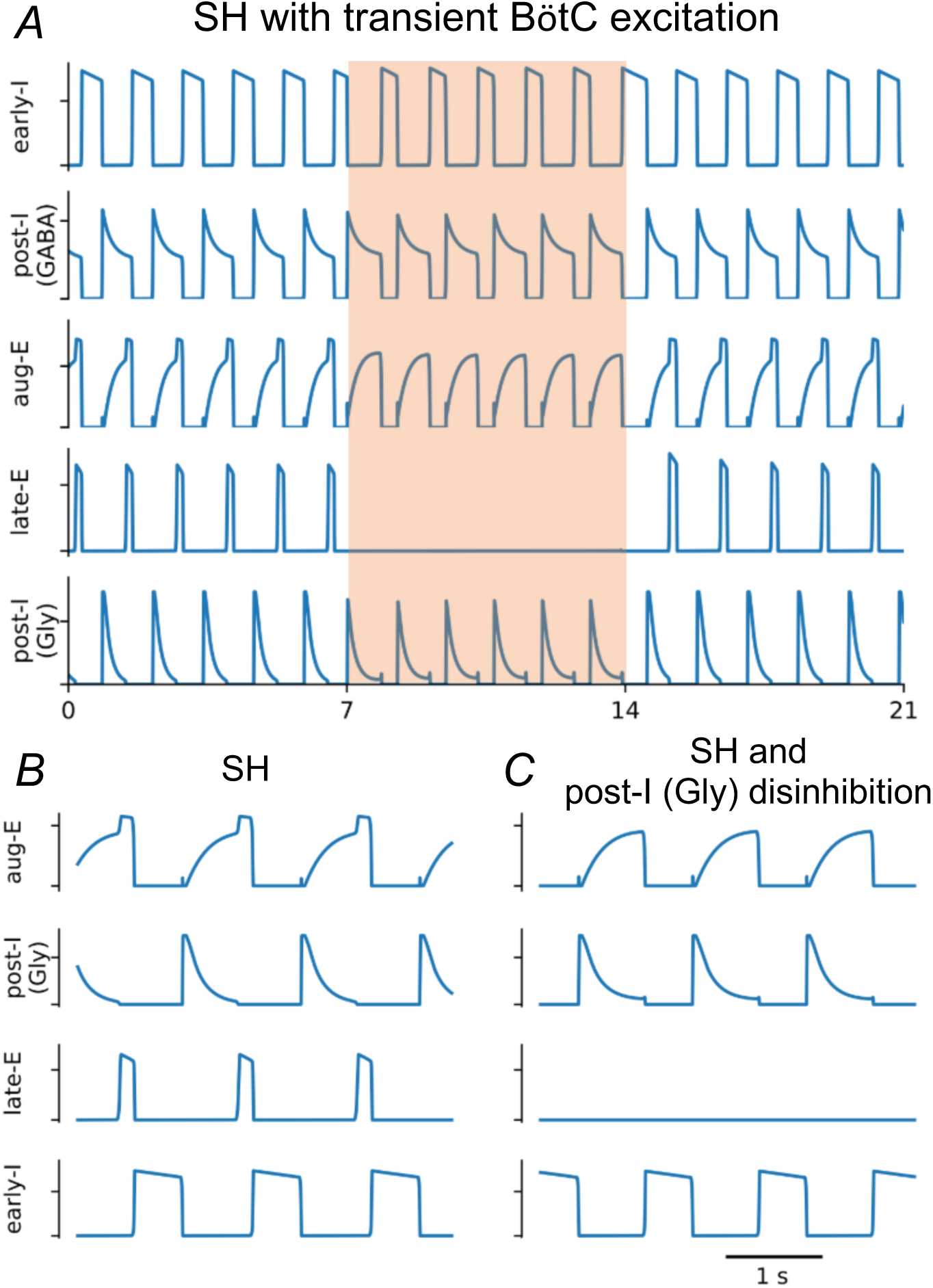
Active expiration induced by reducing drive to neurons active in the post-inspiratory phase was suppressed either by increasing drive to all populations in the Bötzinger complex or by reducing glycinergic transmission within the Bötzinger complex. **Panel A:** Active expiration was induced by decreasing tonic excitatory conductances on the GABAergic (GABA) and glycinergic (Gly) post-inspiratory populations (post-I) by 5%. Active expiration was then abolished by transiently increasing tonic excitatory conductances of the all populations in the Bötzinger complex [post-I and augmenting-expiratory (aug-E)] by 12% during the interval emphasized by the orange shaded box. **Panel B and C:** Active expiration was abolished by reducing the aug-E to post-I synaptic weight by 10%. Other abbreviations – early-I: early inspiratory population, and late-E: late expiratory population.

Taking into account our simulation of hypercapnic conditions, one can also predict that SH-induced active expiration could be abolished by reducing the glycinergic transmission in the BötC, such as achieved following injections of strychnine within this region. We implemented this scenario in the model simulation by reducing the aug-E to post-I_Gly_ synaptic weight by 10%. This change also eliminated active expiration (Figure 10 B and C). The manipulation used in this simulation is identical to the changes implemented in the simulations that depict the reduced glycinergic transmission during the active expiration evoked by high chemosensitive drive. The reduction of the aug-E to post-I_Gly_ synaptic weight disinhibited the post-I_Gly_ population, which resulted in increased inhibition on late-E neurons that were no longer able to fire expiratory bursts (Figure 10 B and C).

### L-glutamate and strychnine microinjections in the Bötzinger complex eliminated the active expiration observed after the exposure to sustained hypoxia

In the next series of experiments, we tested the aforementioned model predictions with microinjections of L-glutamate or strychnine to cause stimulation or disinhibition of BötC of SH rats. We found that L-glutamate microinjections (10 mM) in the BötC of *in situ* preparations of control group (n=7) reduced the PN burst frequency and amplitude, diminished the cVN activity, and produced negligible changes in the AbN activity (Figure 11). On the other hand, in the *in situ* preparations of SH rats (n=7), the response of reduction in PN burst frequency induced by L-glutamate into the BötC was blunted (Δ: −20±9 vs −1±9 bpm, P=0.0019, Figure 11 A and B), while the inhibitory effects on the PN burst amplitude (Δ: −9±6 vs −8±5 % from baseline, P=0.8128, Figure 11C) and cVN activity (Δ: −17±8 vs −13±17 % from baseline, P=0.8048, Figure 11D) were similar to the responses observed in the control group. Moreover, stimulation of the BötC of SH rats markedly reduced the AbN activity due to the transient elimination of late-E bursts during E2 phase (Δ: −4±40 vs −49±30 % from baseline, P=0.0344, Figure 11 A and E). The site of unilateral microinjection in the BötC is illustrated in Figure 11F.

**Figure 11.**
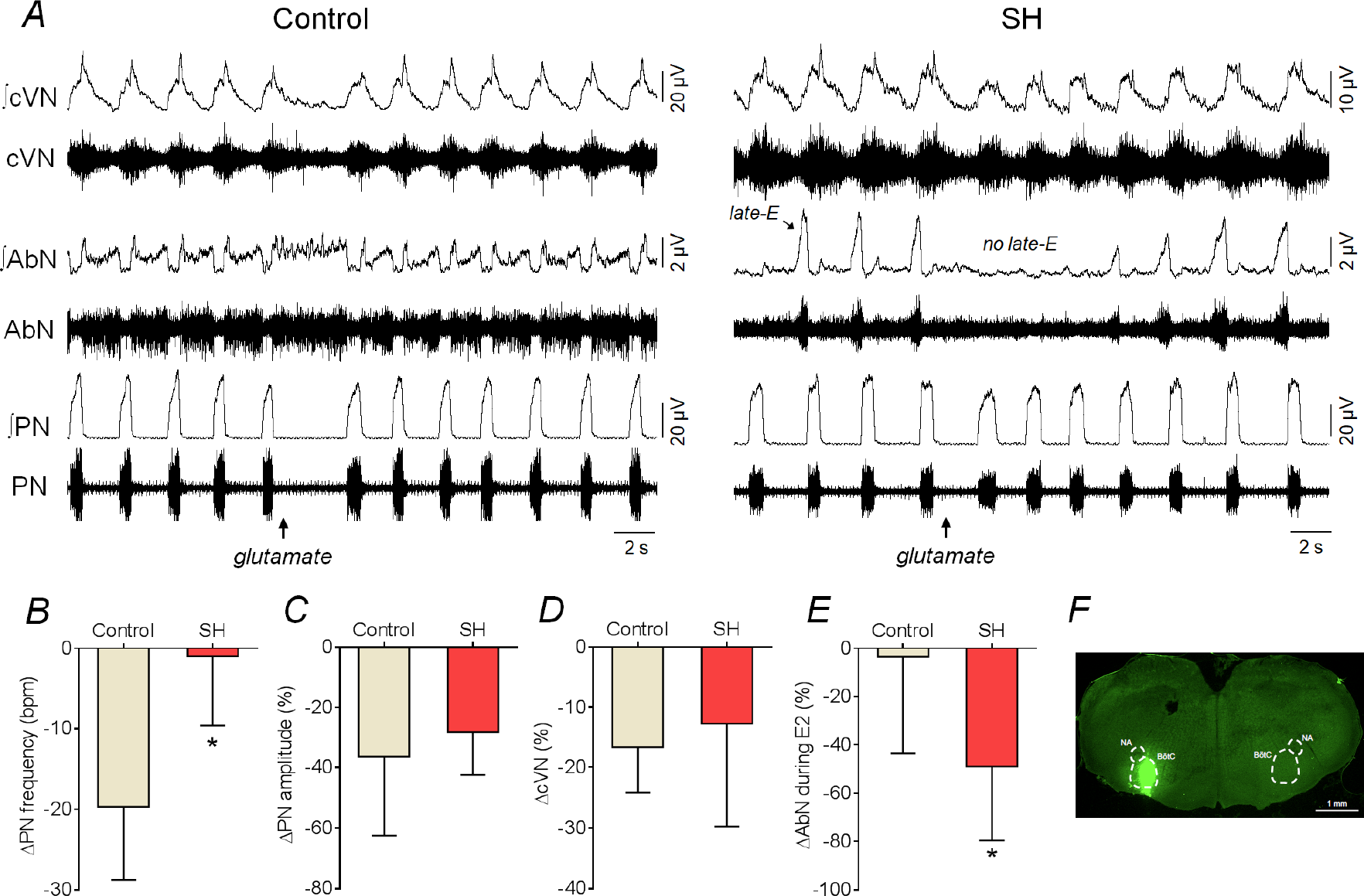
Respiratory motor responses to glutamatergic stimulation of BötC in control and SH rats. **Panel A:** raw and integrated (∫) recordings of cervical vagus (cVN), abdominal (AbN) and phrenic (PN) nerve activities of control and SH *in situ* preparations, representative from their groups, illustrating the respiratory responses to unilateral L-glutamate microinjection (10 mM) in the BötC (arrow). **Panels B-E**: average changes in PN burst frequency and amplitude, cVN post-I and AbN E2 activities, respectively, in response to L-glutamate microinjections in the BötC of control (n=7) and SH (n=7) *in situ* preparations. * different from control group. **Panel F**: coronal section of brainstem from a *in situ* preparation, illustrating the site of unilateral microinjection in the BötC.

As in the experiments with L-glutamate microinjections, disinhibition of the BötC in SH rats with bilateral strychnine microinjections (10 µM) promptly eliminated the pattern of active expiration and restored the eupneic-like respiratory motor pattern in *in situ* preparations (n=8; Figures 12 A and B). After bilateral microinjections of strychnine in the BötC of SH rats, the PN burst frequency increased (15±2 vs 21±7 bpm, P=0.0078, Figure 12C), the PN burst amplitude decreased (41.2±19.3 vs 33.8±19.8 µV, P=0.005, Figure 12D), the time of inspiration increased (0.829±0.197 vs 0.974±0.169 s, P=0.031), the time of expiration reduced (3.407±0.538 vs 2.135±0.611 s, P<0.001), and the cVN post-I mean activity (24.6±4.5 vs 17.9±3.3 µV, P=0.0078, Figure 12E) and duration decreased (73±10 vs 57±12 % of expiratory time, P=0.0007, Figure 12F). Moreover, the antagonism of glycinergic receptors in the BötC of SH reduced the AbN activity during E1 (4.0±2.4 vs 2.6±1.6 µV, P=0.0213, Figure 12G) and E2 phases (8.7±5.1 vs 4.4±2.3 µV, P=0.0072, Figure 12H) – the latter due to the elimination of late-E bursts (Figure 12B). These effects promoted by strychnine in the BötC reversed approximately 30 min after microinjections (data not shown). The sites of bilateral strychnine microinjections in the BötC are illustrated in Figure 12I. Microinjections of vehicle in the BötC of SH rats (n=3) did not change activities of PN, cVN and AbN activities (data not shown). These data indicate that increasing the activity of neurons in the BötC, either by pharmacological stimulation or disinhibition, is able to eliminate active expiration in SH rats.

**Figure 12.**
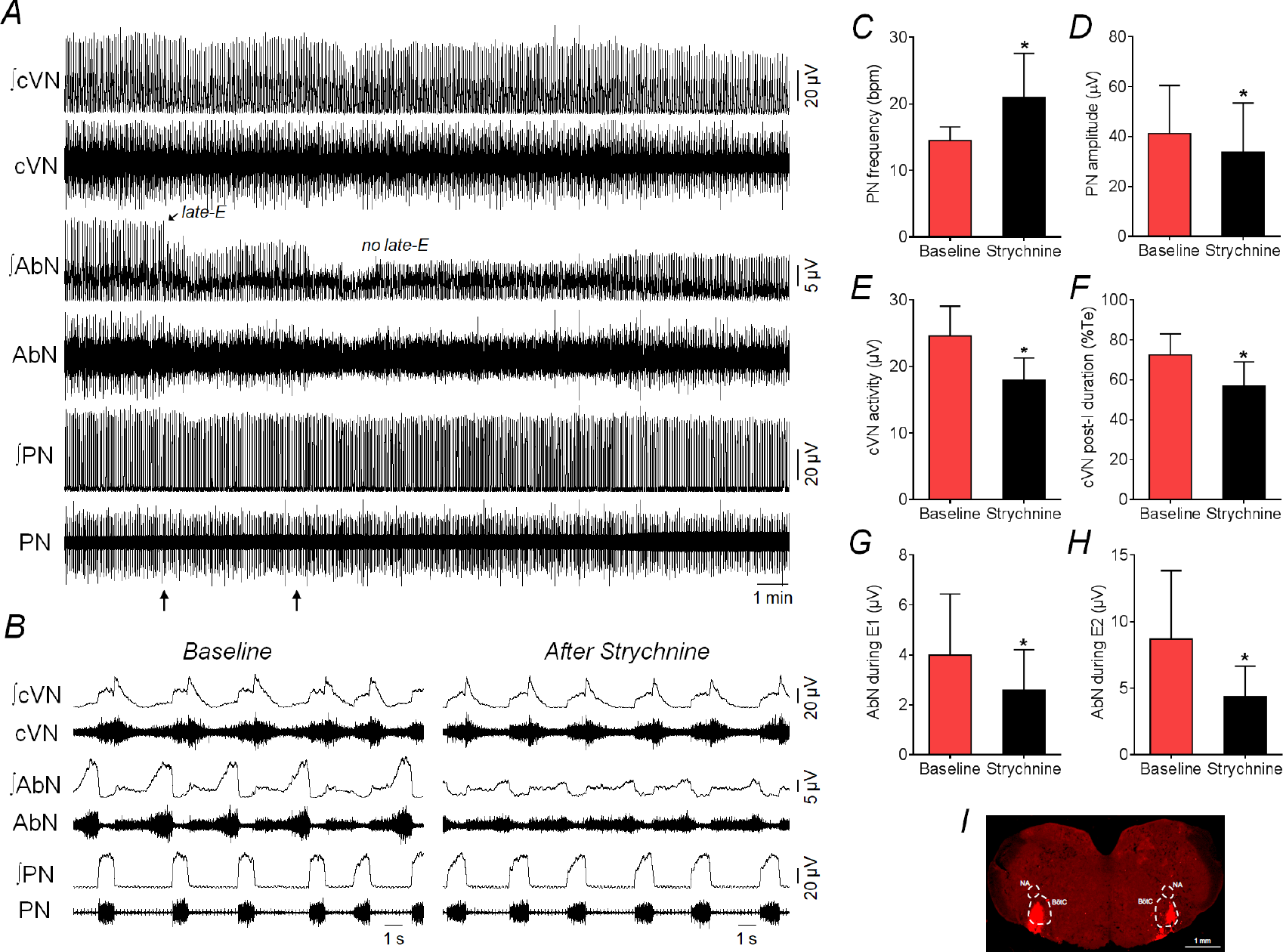
Disinhibition of BötC abolishes active expiration and restores eupneic-like respiratory pattern in SH rats. **Panel A:** raw and integrated (∫) recordings of cervical vagus (cVN), abdominal (AbN) and phrenic (PN) nerve activities of a SH *in situ* preparation, representative from the group, illustrating the effects of bilateral microinjections of strychnine (10 µM) in the BötC (arrows). Note the marked reduction of AbN late-E bursts after the disinhibition of BötC with strychnine. **Panel B**: expanded recordings from panel A, showing, in details, the respiratory motor pattern of SH rats before and after microinjections of strychnine in the BötC. **Panels C-H**: average values of PN burst frequency and amplitude, cVN post-I activity and duration, and AbN activity during E1 and E2 phases, before and after microinjections of strychnine in the BötC of SH *in situ* preparations (n=8). * different from baseline, P<0.05. **Panel I**: coronal section of brainstem from a SH *in situ* preparation, illustrating the sites of bilateral microinjections in the BötC.

### Stimulation of the BötC is sufficient to attenuate the active expiration evoked by glycinergic receptor antagonism in the pFRG in silico and in situ

Our model implies that pFRG late-E activity is restrained by both GABAergic and glycinergic inputs from BötC. However, in our simulations, the partial blockade of glycinergic transmission within the pFRG was sufficient to induce active expiration. We accomplished this by reducing the post-I to late-E synaptic weight by 80% (Figure 13A and B). Therefore, the remaining primarily GABAergic inhibition in this state was not sufficient to prevent late-E neurons from activating at the end of expiration. However, the facilitation of neuronal activity within the BötC, which includes an increase in post-I_GABA_ population activity, could suppress late-E activity induced by reduction of glycinergic transmission within the pFRG. We tested this prediction by increasing the weight of the excitatory drive to each of the three BötC complex populations by 12%, as we previously used in similar simulations for SH condition. The result in simulation was the reduction in frequency of late-E bursts (one late-E burst per two respiratory cycles; Figure 13C) but not the complete elimination of late-E activity.

**Figure 13.**
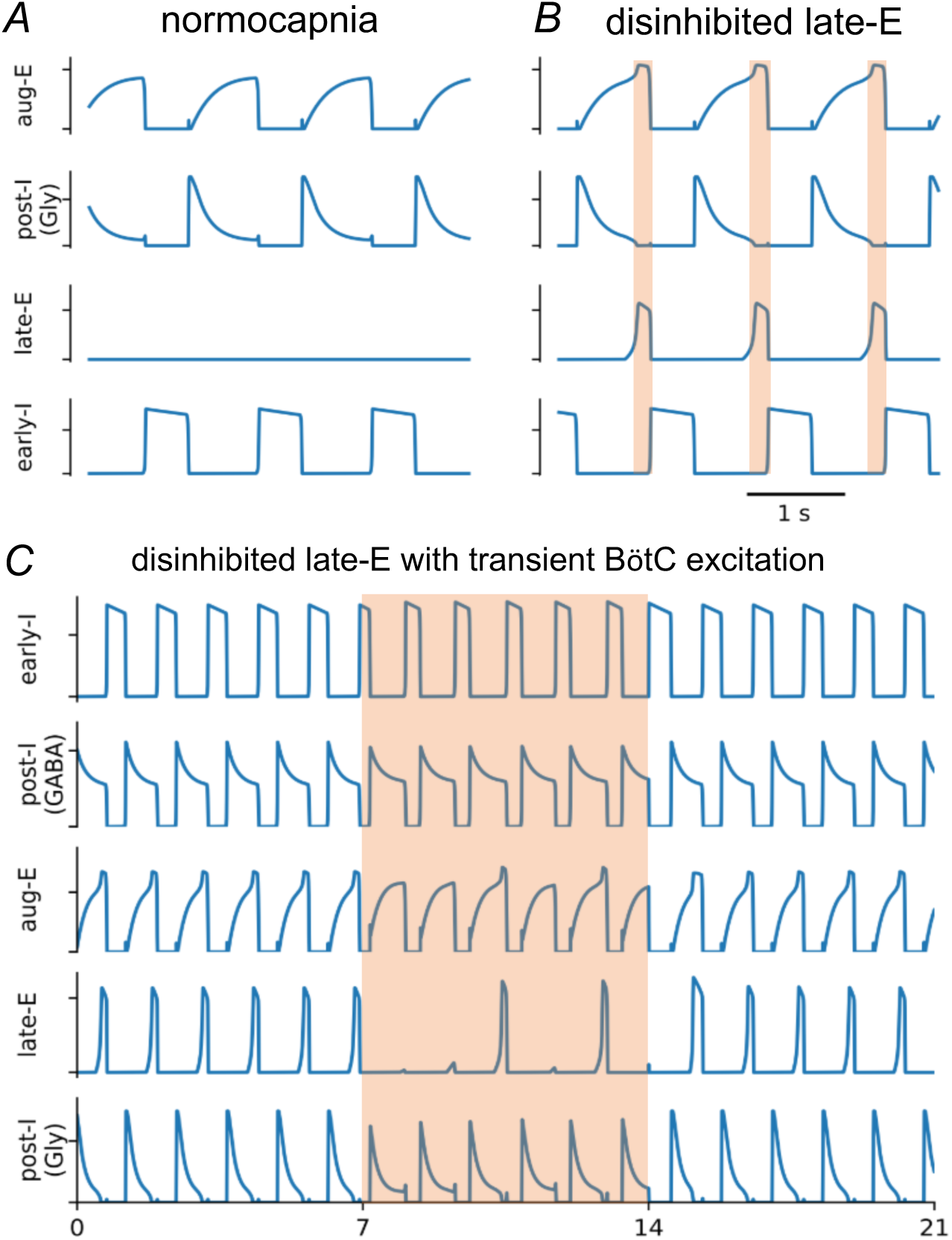
Active expiration was induced by the antagonism of glycinergic receptors within the pFRG, which can be attenuated by an increase in the drive to expiratory populations of the BötC. **Panels A and B:** the reduction of the synaptic projection from the glycinergic post-inspiratory population [post-I (Gly)] to the late-expiratory population (late-E) induced active expiration. The transformation of the respiratory pattern during the late expiratory phase is emphasized by the orange shaded boxes. **Panel C:** In a simulation of this condition, an increase in the tonic excitatory conductances to the populations of the Bötzinger complex — the glycinergic and GABAergic post-I [post-I (GABA)], and the augmenting-expiratory (aug-E) populations – restrained active expiration during the interval emphasized by the orange shaded box.

We then tested this modeling prediction experimentally using *in situ* preparations of control (naïve) rats. Bilateral microinjections of strychnine in the pFRG at 1 mM (but not lower concentrations, data not shown) evoked active expiration in control *in situ* preparations under normocapnia (n=9, Figures 14A), with the emergence of AbN late-E bursts (4.4±2.3 vs 7.1±3.5 µV, P=0.0126, Figure 14C) in all respiratory cycles (Figure 14D). No significant changes were noted in PN burst frequency (21±5 vs 20±4 cpm, P>0.99, Figure 14E) and amplitude (45.4±17.9 vs 43.6±15.3 µV, P0.6961, Figure 14F), as well as in cVN activity (27.1±3.8 vs 26.6±3.6 µV, P>0.99, Figure 14G). Notably, strychnine microinjections in the pFRG was able to attenuate the inhibition of AbN late-E bursts induced by microinjections of L-glutamate (10 mM) in the BötC (Figure 14B), which were observed at similar amplitude (6.0±2.6 µV, P=0.0768, Figure 14B and C) but at lower frequency (12±6 events/min, P=0.0105, Figure 14D) compared to the values observed before L-glutamate microinjections. Moreover, the antagonism of glycinergic receptors in the pFRG prevented the inhibitory effects on PN burst frequency (21±4 cpm, P=0.1676, Figure 14E), but not on PN burst amplitude (38.2±14.1 µV, P=0.0294, Figure 14F) and cVN activity (21.3±2.6 µV, P=0.0057, Figure 14 G) induced by glutamatergic stimulation of the BötC. The sites of microinjections within the pFRG and in the BötC are illustrated in Figure 14H. These results agree with modeling predictions, indicating that glycine inputs are critical to restrain the activity of pFRG oscillatory under normocapnic conditions, and that stimulation of BötC neurons after the antagonism of glycine receptors in the pFRG partially attenuates the emergence of late-E activity.

**Figure 14.**
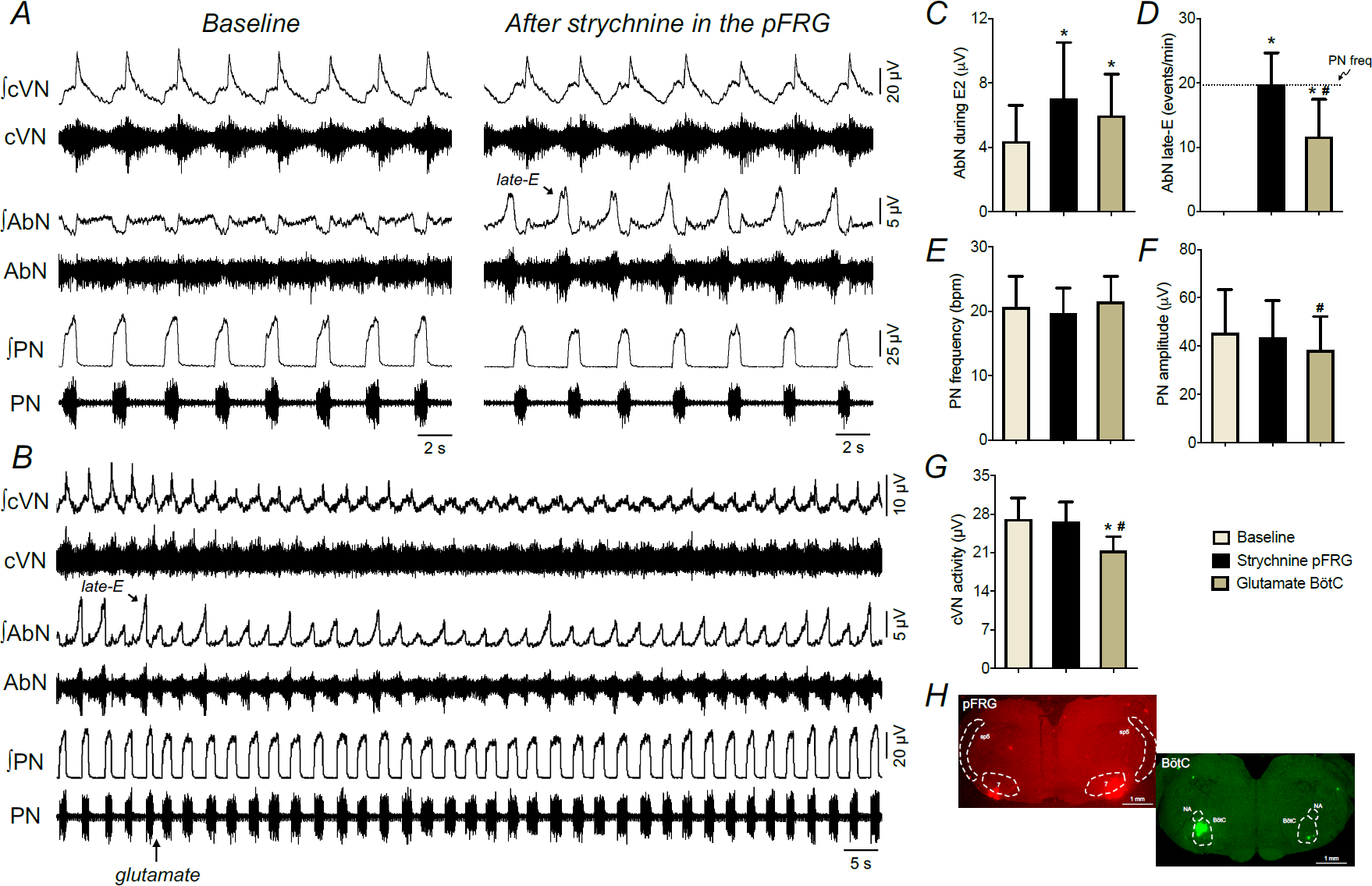
The antagonism of glycine receptors in the pFRG attenuated the inhibitory effect of BötC stimulation on the active expiratory pattern in control rats. **Panel A:** raw and integrated (∫) recordings of cervical vagus (cVN), abdominal (AbN) and phrenic (PN) nerve activities of a control *in situ* preparation, representative from the group, before and after bilateral microinjections of strychnine (1 mM) in the pFRG. Note the generation of active expiration (AbN late-E bursts) after disinhibition of pFRG. **Panel B**: recordings of cVN, AbN and PN of a control *in situ* preparation, illustrating the respiratory responses to unilateral L-glutamate microinjection (10 mM) in the BötC (arrow) after the antagonism of glycine receptors in the pFRG. **Panels C-G**: average values of PN burst frequency and amplitude, cVN post-I and AbN E2 activities, and AbN late-E frequency, respectively, under baseline conditions, after strychnine microinjections in the pFRG and after L-glutamate microinjections in the BötC of control *in situ* preparations (n=9). * different from baseline, # - different from strychnine in the pFRG, P<0.05. **Panel H**: coronal sections of brainstem from a control *in situ* preparation, illustrating the sites of bilateral microinjections in the pFRG (red) and unilateral microinjection in the BötC (green).

## DISCUSSION

Contractions of pump abdominal muscles are observed in conditions of high metabolic demand, and are an important mechanism to improve pulmonary ventilation and gas exchange (Jenkin and Milsom, 2014, Lemes and Zoccal, 2014). This active expiratory pattern emerges from to the stimulation of the conditional expiratory oscillatory located in the pFRG (Janczewski and Feldman, 2006). In addition to excitatory mechanisms (Abdala et al., 2009, Zoccal et al., 2018), inhibition to the pFRG has also been suggested to control the recruitment and timing of abdominal bursts within the expiratory phase (Pagliardini et al., 2011, de Britto and Moraes, 2017, Barnett et al., 2018, Molkov et al., 2010). Here, we present new experimental evidence indicating that the BötC is an important source of inhibitory inputs that restrain the occurrence of active expiration. Combining *in situ* and *in silico* approaches, we propose that the BötC and the pFRG establish dynamically bidirectional interactions, where the post-I neurons of the BötC provide inhibition to the pFRG peaking at early expiration, while the pFRG neurons, when recruited, excite BötC aug-E neurons during the late expiration. Moreover, our data further suggest a specific inhibitory connectome among the BötC neurons that is consistent with differential effects of manipulations with GABAergic and glycinergic neurotransmission within the BötC on active expiration. This study advances our understanding of the neural components and synaptic inputs that provide neural control of the active expiratory pattern in rats.

Previous evidence indicates that excitatory and inhibitory synapses are important to recruit and shape the activity of the late-expiratory population of the pFRG and, hence, contribute to the emergence of active expiration (Molkov et al., 2010). Using mathematical models, we previously proposed that strong inhibition during inspiration and early expiration prevents the activation of the late-E population of the pFRG under resting conditions. However, as inhibition during E2 phase decreases, late-E neurons may overcome the expiratory inhibition in the presence of enhanced excitatory drive, and fire action potentials (Molkov et al., 2010, Rubin et al., 2011). Experimentally, the pre-BötC and the RTN neurons have been suggested as important sources of excitation to the pFRG (Huckstepp et al., 2016, Zoccal et al., 2018). On the other hand, the sources of inhibitory inputs to the pFRG still remain to be elucidated. Our findings support the notion that the BötC provides functional inhibitory inputs to the pFRG. The BötC contains inhibitory neurons that are most active during the E1 (post-I population) and E2 phases (aug-E population) and critically control the transition from inspiration to expiration in eupnea (Smith et al., 2007). During hypercapnia, we found that both neuronal populations showed an increase in their average discharge frequency. On the other hand, the association between active expiration and firing pattern differed between post-I and aug-E neurons. Regarding to the aug-E population, we found that these neurons were stimulated in the presence of active expiration. Our findings parallels with the results by Abdala et al. (2009) that demonstrated that the late expiratory activation of aug-E neurons during hypercapnia can be blocked by the pharmacological inhibition of the RTN/pFRG neurons. Altogether, these observations indicate that excitatory inputs originating from the pFRG or RTN during hypercapnia/active expiration excite BötC aug-E neurons during the E2 phase. This might represent a mechanism that delays of onset of inspiration and prolongs the expiratory duration when abdominal late-E bursts occur (Abdala et al., 2009, Molkov et al., 2010, Zoccal et al., 2018). Studies by Abdala et al. (2009) also proposed an inverse association between the BötC post-I activity and active expiration, based on the observation that vagal post-inspiratory motor activity reduces when abdominal late-E bursts emerge during hypercapnia. Herein, we provide direct and assertive evidence demonstrating that post-I neurons of the BötC stop firing during strong abdominal late-E activity evoked by hypercapnia. These findings show that although both populations receive excitatory inputs during hypercapnia, possibly from CO_2_-chemosensitive sites, post-I and aug-E neurons of the BötC are distinctly controlled by the network responsible for the generation of active expiration.

The experiments with pharmacological manipulations of the BötC were then performed to gain information about the functionality of BötC neurons during active expiration. Based on the fact that neurons that express GAD67 and GlyT2 mRNA are found in the BötC region (Tanaka et al., 2003), and on the fact that GABAergic and glycinergic synapses are utilized by the BötC neurons during eupnea (Ezure et al., 2003a, Ramirez et al., 1997), we promoted the disinhibition of the BötC utilizing either glycinergic or GABA_A_ receptor antagonists, strychnine and gabazine, respectively. It is important to mention that the concentrations of strychnine and gabazine used in our study did not disrupt the 3-phase respiratory pattern. Surprisingly, these antagonisms generated opposite effects: while glycine within the BötC was required for the abdominal late-E breakthrough during hypercapnia (strychnine microinjections attenuated the abdominal late-E bursts), GABAergic neurotransmission in the BötC was necessary to restrain active expiration at rest (gabazine microinjections generated active expiration under normocapnic conditions). These data therefore indicate that the inhibitory neurons of the BötC may form specific patterns of synaptic connections, using distinctly glycine and GABA, to control abdominal expiratory activity.

Using our theoretical model of respiratory network in combination with these new experimental data, we generated hypotheses about the role of BötC neurons for the inhibitory control of the expiratory motor pattern formation. In this model, we proposed two different types of inhibitory post-inspiratory cells, one that utilizes GABA as the neurotransmitter (named post-I_GABA_) and another that uses glycine (named post-I_Gly_). We also assumed that the post-I_GABA_ cells sent GABAergic inputs to the aug-E neurons, while the post-I_Gly_ neurons received glycinergic inputs from the aug-E population. Moreover, both GABAergic and glycinergic post-I populations sent inhibitory projections to the pFRG neurons. With this framework, our model generated responses in simulated conditions qualitatively equivalent to the experimental data. We observed that the emergence of active expiration during hypercapnia required an increased excitatory drive from chemosensitive sites to the pFRG, as previously demonstrated (Zoccal et al., 2018). The activation of the late-E neurons in the pFRG promoted the stimulation of aug-E neurons in the BötC that, in turn, depressed the activity of pFRG-projecting glycinergic post-I neurons, facilitating the late-E breakthrough during E2 phase. According to the model, the ablation of active expiration caused by strychnine microinjections in the BötC resulted from the disinhibition of post-I_Gly_ neurons, increasing the glycinergic drive to the pFRG neurons. On the other hand, the emergence of active expiration induced by gabazine injections in the BötC during normocapnia was consequent to the attenuation of GABAergic inputs on the aug-E neurons, that ultimately increased the inhibitory drive on the post-I_Gly_ neurons, disinhibiting the pFRG expiratory neurons. We assumed in our model that GABAergic inputs originated within the BötC, but we acknowledge that GABAergic inputs may also originate from other regions such as pre-BötC, NTS and Raphe (Kubin et al., 2006, Iceman and Harris, 2014, Marchenko et al., 2016). Although these hypotheses are theoretical and still required appropriated experiments to be proven, the model suggests a novel possible synaptic organization of the BötC inhibitory circuit, with differential role for inhibitory synapses (GABA vs glycine) across different states (normocapnia vs hypercapnia) to control active expiration.

To experimentally test the model predictions about the role of the BötC neurons during active expiration, we also performed experiments on rats exposed to short-term sustained hypoxia. This is an attractive model to investigate the neuronal circuitry and dynamics responsible for the generation of active expiration because it causes a long-lasting activation of the pFRG expiratory neurons under resting conditions (Moraes et al., 2014). We previously demonstrated that rats exposed to 24 h of 10% O_2_ exhibited a persistent increase in minute ventilation upon reoxygenation [named ventilatory acclimatization to hypoxia, VAH, (Powell et al., 1998)] that was associated with the emergence of active expiratory pattern, mediated by plastic changes in central mechanisms (Flor et al., 2018). Therefore, the use of SH-treated rats to verify modeling predictions would also contribute to expand our understanding about the mechanisms associated with the development of VAH. In agreement with previous studies (Moraes et al., 2014), we found that the BötC post-I neurons of SH preparations were silent during the emergence of late-E activity, while aug-E neurons were excited in the presence of active expiation – discharge patterns that resemble the results observed in naïve rats exposed to hypercapnia. In the model, the ‘SH condition” was simulated by reducing the excitatory drive to the GABAergic and glycinergic post-I neurons. This hypothesis is in agreement with previous observations that the depressed the activity of post-I neurons in SH is synaptically mediated (Moraes et al., 2014). In this scenario, we consider that changes in the pontine excitatory drive to the VRC, specially from the Kölliker-Fuse (Dutschmann and Herbert, 2006, Geerling et al., 2017, Barnett et al., 2018, Jenkin et al., 2017), or reductions in the excitatory drive from the recently described post-inspiratory complex (Anderson et al., 2016), may represent possible mechanisms underlying the depressed activity of post-I neurons in the BötC after SH – hypotheses that still require additional studies to be addressed.

Using the SH condition in this model, we predicted that manipulations that increase the activity of post-I neurons would be able to abolish the active expiration in SH-treated rats. Specifically, our simulations demonstrated that either a general stimulation of the BötC neurons, or a selective increase in the activity of post-I glycinergic neurons would be sufficient to eliminate the late-E activity in the abdominal motor output of SH-conditioned rats. The experimental data confirmed the modeling predictions. In agreement with previous experimental studies (Moraes et al., 2012b, Moraes et al., 2011), unilateral microinjection of glutamate in the BötC produced a transient increase in expiratory duration (hence decreasing the respiratory frequency) and reduced phrenic activity in *in situ* preparations of control rats, possibly due to the inhibition of inspiratory neurons of the pre-BötC/rVRG (Ezure et al., 2003c, Smith et al., 2007). We also noticed that glutamate in the BötC depressed central vagus activity – a response that may have been caused by the stimulation of inhibitory neurons that control pre-motor/motor neurons associated with the formation of vagal motor activity (Smith et al., 2007). Moreover, stimulation of BötC with glutamate of control rats did not modify resting abdominal activity. In SH rats, unilateral glutamate microinjections in the BötC did not promote changes in the respiratory frequency but reduced the phrenic burst and central vagus activity, as well as eliminated the late-E bursts in abdominal. The transient cessation of late-E abdominal activity in SH rats is consistent with the notion that glutamate microinjection stimulated BötC inhibitory neurons that suppressed pFRG neurons. After bilateral microinjections of strychnine in the BötC of SH rats, we observed a long-lasting suppression of abdominal late-E activity, and the recovery of the resting 3-phase eupneic respiratory pattern – an effect not observed in control animals. These findings strongly support our hypothesis that BötC neurons that are under the control of glycinergic neurotransmission, supposedly the post-I population, are critical for the control of abdominal expiratory activity, so that a reduction in their baseline activity can lead to the emergence of active expiration in rats exposed to SH.

Studies demonstrating that the antagonism of glycinergic and GABAergic receptors in the pFRG evokes active expiration (Pagliardini et al., 2011, de Britto and Moraes, 2017) indicate that the expiratory neurons of the pFRG are synaptically supressed under conditions of normoxia/normocapnia. According to our data, part of these inhibitory projections originates from the BötC. It was previously shown that active expiration can be evoked by blocking GABAergic inhibition only within the pFRG (Molkov et al., 2010). This was considered in our model by the GABAergic projections from the post-I_GABA_ population of the BötC and early-I population in the pre-BötC to the late-E neurons in the pFRG. However, our model also suggested a relevant role for the glycinergic projections from BötC post-I neurons to the pFRG, as the suppression of this pathway generated active expiration under normocapnic conditions. This prediction was verified experimentally, as bilateral microinjections of strychnine in the pFRG not only evoked active expiration, but also greatly attenuated the inhibitory effect on the abdominal activity elicited by microinjections of glutamate in the BötC. The residual abdominal inhibitory response elicited by glutamate in the BötC after strychnine in the pFRG may be elicited by the GABAergic projections from the BötC to the pFRG, as suggested by the model. We cannot exclude the possibility that our microinjections in the pFRG did not block all glycine receptors. After microinjections of strychnine in the pFRG, as well as in animals exposed to SH, the microinjections of glutamate in the BötC did not prolong expiratory duration and did not reduce the respiratory frequency. These findings suggest that the recruitment of pFRG expiratory neurons may also modify the activity/dynamics of BötC neurons that inhibit the inspiratory neurons. Alternatively, we may also consider that the possible interactions between the pFRG neurons and the inspiratory rhythm generating neurons in the pre-BötC (Huckstepp et al., 2016) are modifying the pattern of response to glutamate in the BötC. Additional studies are required to verify these possibilities.

In conclusion, our study provides evidence supporting the notion that the BötC neurons are an essential part of the neural circuitry controlling the generation of active expiration. Our data provide further mechanistic details for our previous studies indicating that the neurons of the Kölliker-Fuse, in the dorsolateral pons, coordinate the transition from post-I to late-E phases and control the onset of abdominal late-E busts (Barnett et al., 2018, Jenkin et al., 2017). Altogether, these findings indicate that a pontine-medullary circuitry, through connections from Kölliker-Fuse to the BötC, is required for the inhibitory control of pFRG, critically contributing for the formation of abdominal expiratory activity. Neuroplasticity in this pathway may be responsible for the changes in breathing following the exposure to chronic challenges, such as sustained hypoxia (as experienced in high altitudes), or contribute to pathological respiratory phenotypes, as seen in sleep breathing disorders, heart failure, neurological diseases (e.g. Rett syndrome) and others.

## MATERIALS AND METHODS

### Ethical approval

Experiments were performed on juvenile male Holtzman rats (60 – 80 g, 23 – 28 days old) obtained from the Animal Care Unit of the São Paulo State University, Araraquara. The animals were housed in collective cages with free access to rat chow and water, under controlled conditions of temperature (22 ± 1 °C), humidity (50 - 60%) and light/dark cycle (12:12 lights on at 07:00 am). All experimental procedures comply with the Guide for the Care and Use of Laboratory Animals published by the Brazilian National Council for Animal Experimentation Control (CONCEA), conform to the principles and regulations under which the journal operates, and were approved by the Local Ethical Committee in Animal Experimentation of School of Dentistry of Araraquara, São Paulo State University (protocol 30/2016). All steps were taken to avoid animals’ pain and suffering.

### Sustained Hypoxia

One day before the experiments, animals were housed in collective cages and placed inside chambers that allowed the control of the inspired oxygen fraction via injections of pure N_2_ and O_2_ (White Martins, Sertãozinho, Brazil) through a computerized system of solenoid valves (Oxycycler, Biospherix, Lacona, NY, USA). The conditions of the temperature, humidity, light/dark cycle and food/water access inside the chambers were kept in standard conditions as aforementioned. In the hypoxic chamber, the animals were exposed to 10% O_2_ for 24 h (sustained hypoxia, SH); whilst in the control chamber the animals were maintained under normoxic condition (Flor et al., 2018, Moraes et al., 2014). The levels of O_2_ inside the chambers were monitored continuously and the gas injections were performed in the upper portion of the chambers to prevent air jets from directly reaching the animals. Experiments were performed just after the end of the experimental protocols.

### In situ arterially perfused rat preparations

The animals were surgically prepared to obtain the *in situ* working heart-brainstem preparations as previously described (Zoccal et al., 2008, Flor et al., 2018). Briefly, animals were heparinized (1,000 IU) and subsequently deeply anaesthetized with isoflurane until the paw and tail pinch reflexes were abolished, then transected sub-diaphragmatically, exsanguinated (that resulted in euthanasia) and immersed in cold (2-4° C) Ringer solution (in mM: 125 NaCl; 25 NaHCO_3_; 3.75 KCl; 2.5 CaCl_2_; 1.25 MgSO_4_; 1.25 KH_2_PO_4_ and 9.9 glucose). The preparations were then decerebrated at precollicular level and skinned, the lungs were removed, and descending aorta was isolated. After, the preparations were placed in supine position, and the trachea, esophagus and all muscles and connective tissues covering the occipital bone were removed. The basilar portion of the atlanto-occipital membrane was cut, and the occipital bone was carefully removed using a micro-Rongeur (Fine Scientific Instruments, England) to expose the ventral surface of the medulla, from the vertebral arteries to the beginning of the ventral surface of the pons. Subsequently, the preparations were transferred to a recording chamber, the descendent aorta was isolated, cannulated with a double-lumen cannula and perfused retrogradely (21 - 25 mL.min^−1^; Watson-Marlow 520S, Falmouth, UK) with modified Ringer solution containing 1.25% polyethylene glycol 20,000 (oncotic agent; Sigma-Aldrich, USA) and vecuronium bromide (neuromuscular transmission blocker; 3 - 4 μg.mL^−1^). The perfusion pressure was held between 55-75 mmHg by the addition of vasopressin (0.6 - 1 nM, Sigma, USA) to the perfusate. The perfusion solution was constantly gassed with 95% O_2_ - 5% CO_2_ (pH 7.4), warmed to 31 - 32°C and filtered using a polypropylene filter (25 μm pore size). The perfusion pressure was recorded with a pressure transducer (MLT06070, ADInstruments, Bella Vista, Australia) connected to an amplifier (Grass Quad Amplifier, model 15LT, RI, USA).

### Nerve recordings and analyses

Respiratory motor nerves were isolated and recorded simultaneously using bipolar suction electrodes held in micromanipulators (Narishige, Tokyo, Japan). To measure the inspiratory motor output, the left phrenic nerve (PN) was isolated and cut distally at its insertion to the diaphragm. Its rhythmic ramping activity provided a continuous physiological index of preparation variability (Paton, 1996). The motor activity to laryngeal abductor and adductor muscles was evaluated by recordings from the left vagus nerve (cVN), which was dissected and sectioned at the cervical level (below the bifurcation of the common carotid artery). To measure the motor output to abdominal muscles, nerves from the right lumbar plexus at the thoracic-lumbar level (T_12_ - L_1_) were isolated and cut distally and referred to as abdominal nerve (AbN). Bioelectric signals were amplified (CP511 Amplifier, Grass Technologies, RI, USA), filtered (0.3 - 3 kHz) and acquired by A/D converter (micro1401, Cambridge Electronic Design Limited, Cambridge, England) on a computer using Spike 2 software (version 9, Cambridge Electronic Design).

Analyzes of the activities of the recorded nerves were carried out on rectified and smoothed signals (time constant of 50 ms), after the electrical noise subtraction, using custom-written scripts in Spike 2 software, as previously described (Barnett et al., 2018). Using the PN and cVN recordings as reference, the respiratory cycle was divided into 3 phases: i) inspiration, coincident with PN burst; ii) post-inspiration (E1), coincident with the decrementing activity of cVN during the expiratory phase, corresponding to approximately 2/3 of the initial expiratory cycle; and iii) E2, corresponding to the silencing period of the cVN, representing approximately the final third of the expiration. PN activity was evaluated by its burst frequency and amplitude (expressed in bursts per minute, bpm, and µV, respectively). From the PN activity, the duration of inspiratory (PN burst length) and expiratory phases (time interval between consecutive PN bursts) were also determined. The activity of the cVN was quantified as the mean activity (expressed in µV) and duration (expressed as a percentage of the total expiratory time) of the post-inspiratory phase. The mean AbN activity (expressed in µV) was quantified during E1 and E2 phases.

### Single-unit neuronal recordings and analyses

Unitary extracellular recordings of the activity of post-I and aug-E neurons of the BötC were performed in *in situ* preparations as described previously (St-John et al., 2009, Moraes et al., 2012b). Glass microelectrodes (15-35 MΩ, filled with 1M NaCl and 1% BAPTA AM, Sigma-Aldrich) were mounted in a 3D micromanipulator (Scientifica Ltd, Uckfield, UK) and positioned in the BötC with the aid of surgical microscope, considering the anatomical references of the ventral surface of medulla and the following stereotactic parameters: 700 - 900 µm caudal to caudal pole of the trapezoid body; 1500 - 1800 µm lateral to midline and 350 - 500 µm beneath the ventral surface. The signals were amplified (Duo 773 Electrometer, WPI, USA), low pass filtered (filter 2 KHz) and acquired in a microcomputer (micro1401, CED, England) using appropriate software (Spike 2, CED, England). Post-I and aug-E neurons were identified according to their typical pattern of discharge (Smith et al., 2007, Moraes et al., 2014). The neuronal activities were evaluated according to: i) average instantaneous firing frequency (expressed in Hz), calculated from the time interval between consecutive action potential (excluding intervals smaller than 1 ms); and ii) the duration (expressed in seconds) of active and silent (no action potentials) periods during the expiratory phase.

### Exposure to hypercapnia

*In situ* preparations of control rats were exposed to periods of increased fractional concentration of CO_2_ in the perfusate from 5 to 8% CO_2_ (balanced in O_2_) for 5 minutes, using a gas mixer (AVS Projetos, São Carlos, Brazil) connected to an O_2_ and CO_2_ analyzer (ADInstruments, Bella Vista, Australia), to generate the pattern of active expiration (Zoccal et al., 2018). The effects peaked and reached a steady state after 3 min of exposure. The respiratory responses to hypercapnia were quantified as the maximal variations relative to respective baseline activity. A recovery period of at least 10 minutes was considered between consecutive stimuli.

### Intraparenchymal injections in the BötC and pFRG

Borosilicate pipettes (internal diameter of 0.84 mm; Sutter Instruments, USA), were pulled and used for intraparenchymal injections in the BötC and the pFRG. The micropipettes were connected to a pneumatic pump (Picospritzer II - Parker Hannifin Corporation, Fairfield, USA), which allowed the microinjection of the drugs by means of calibrated “pulses of pressure” to a volume of approximately 35 - 40 nL. This apparatus was then adapted in a high-resolution 3D micromanipulator (Narishige, Japan), and then the micropipettes were positioned, with the aid of a surgical microscope, in the areas of interested using the following stereotaxic coordinates (Moraes et al., 2012a, Moraes et al., 2012b): i) BötC: 700 - 900 µm caudal to caudal pole of the trapezoid body, 1500 - 1800 µm lateral to midline and 350 - 500 µm beneath the ventral surface; and ii) pFRG: 400 to 500 µm caudal to the caudal pole of the trapezoid body; 1800 to 2000 µm lateral to midline; and 40 to 50 µm beneath the ventral surface. L-glutamate (10 mM, excitatory amino acid) was microinjected unilaterally whilst strychnine (10 µM in the BötC and 1 mM in the pFRG, glycine receptor antagonist) and gabazine (250 µM, GABA_A_ receptor antagonist) was microinjected bilaterally. Bilateral microinjections were completed in a time interval lower than 1 minute (counting from the first microinjection). The drugs were dissolved in Ringer solution containing either green or red fluorescent microspheres (1%, Lummafluor, USA).

### Histology

At the end of experiments involving microinjections in the BötC and pFRG, the brainstems of the *in situ* preparations were rapidly removed and immersed in a solution containing 10% formaldehyde for 5 days, and then in a solution containing 20% sucrose. Using a cryostat (Leica CM1850, Wetzlar, Germany), sequential coronal sections (50 µm thickness) containing the pFRG, BötC and adjacent regions were obtained and analyzed in a fluorescence microscope (Leica DM IRB, Wetzlar, Germany). The sites of L-glutamate and gabazine microinjections were identified by visualization of green microspheres whilst the sites of strychnine or vehicle (Ringer) microinjections were identified by the visualization of red microspheres.

### Data analyses

The results were described and graphically represented as mean ± standard deviation (SD). The normal distribution of the data was verified with Shapiro-Wilk normality test. With respect to neuronal activity of control rats, paired Student’s t-test and two-way ANOVA followed by the post-test of Bonferroni was used for the analyses of firing frequency and duration of active/silent periods, respectively. The respiratory responses to unilateral L-glutamate microinjections, as well as the comparisons of respiratory parameters between control and SH rats, were analyzed using unpaired Student’s t-test. The respiratory responses to bilateral microinjections of strychnine in the BötC, and strychnine microinjections in the pFRG followed by microinjections of L-glutamate in the BötC were analyzed using repeated measurements one-way ANOVA followed by post-test of Bonferroni. Significant differences were considered when P <0.05. Statistical and graphic operations were performed using GraphPad Prism software (version 8, GraphPad, La Jolla, USA).

### Mathematical modeling

We have extended our previous firing-rate based model of the respiratory central pattern generator to account for differential respiratory responses to perturbations of glycinergic and GABAergic transmission within the BötC (Figure 5). The respiratory circuitry in this model was based on the respiratory circuitry presented in a series of previous models that examined the emergence of active expiration as well as gas-exchange and pulmonary feedback (Rubin et al., 2009, Rubin et al., 2011, Molkov et al., 2014, Molkov et al., 2016). The network connectivity in these models was informed by brainstem transection experiments presented in Smith et al. (2007). The three phase respiratory rhythm was produced by a circuit that includes the pre-inspiratory/inspiratory (pre-I/I) population of the pre-BötC, which acts as the excitatory kernel of the respiratory central pattern generator (rCPG), and inhibition among the early-inspiratory (early-I) population of the pre-BötC, the augmenting-expiratory (aug-E) population of the BötC, and an additional population within the BötC that fires strongly during the post-inspiratory phase and adapts to a steady state firing rate by the end of expiration.

The principal change implemented in this new version of the model is the discrimination of glycinergic and GABAergic neuronal populations within the BötC. We assumed that the aug-E population is glycinergic. The BötC population that is active during the post-inspiratory phase was split into a GABAergic (post-I_GABA_) and a glycinergic (post-I_Gly_) populations. The post-I_GABA_ population participated in rhythm generation and the formation of the three-phase respiratory pattern. It inhibited aug-E and receives inhibition from early-I populations. Its firing pattern was strong at the beginning of expiration, and it persisted to the E-to-I phase transition. This GABAergic population was also involved in control of active expiration; it inhibited the late expiratory (late-E) population of the pFRG. The new glycinergic post-I population also fired strongly at the beginning of expiration, but its pattern was substantially shaped by inhibition from aug-E. As the firing rate of aug-E ramps during expiration, post-I_Gly_ activity was suppressed. Depending on the physiological constraints imposed upon model simulations, the post-I_Gly_ population fired at a low rate at the end of expiration or was suppressed entirely before the E-to-I transition. This glycinergic population was also involved in the control of active expiration by way of an inhibitory projection to late-E in the pFRG. In this way, we distinguished glycinergic and GABAergic inhibitory projections within the BötC and from the BötC to the pFRG.

We tested the model by simulating the responses of the respiratory CPG to a number of pharmacological and physiological manipulations. The antagonism of GABAergic and glycinergic receptors in the BötC or the pFRG was modeled as a reduction in the weight of synaptic conductances within specific compartments. The response to hypercapnia was modeled as an increase in the chemosensitive drive to the late-E population of the pFRG. The transformation of the respiratory output following SH conditioning was modeled by reducing the conductance of the tonic excitatory drives on the GABAergic and glycinergic post-I populations (SH modulated drive). The injection of glutamate specifically to the BötC was modeled as a uniform increase in the tonic excitatory drive to all of the BötC populations. The precise distribution and extent of these perturbations are summarized in Table 1.

**Table 1.**
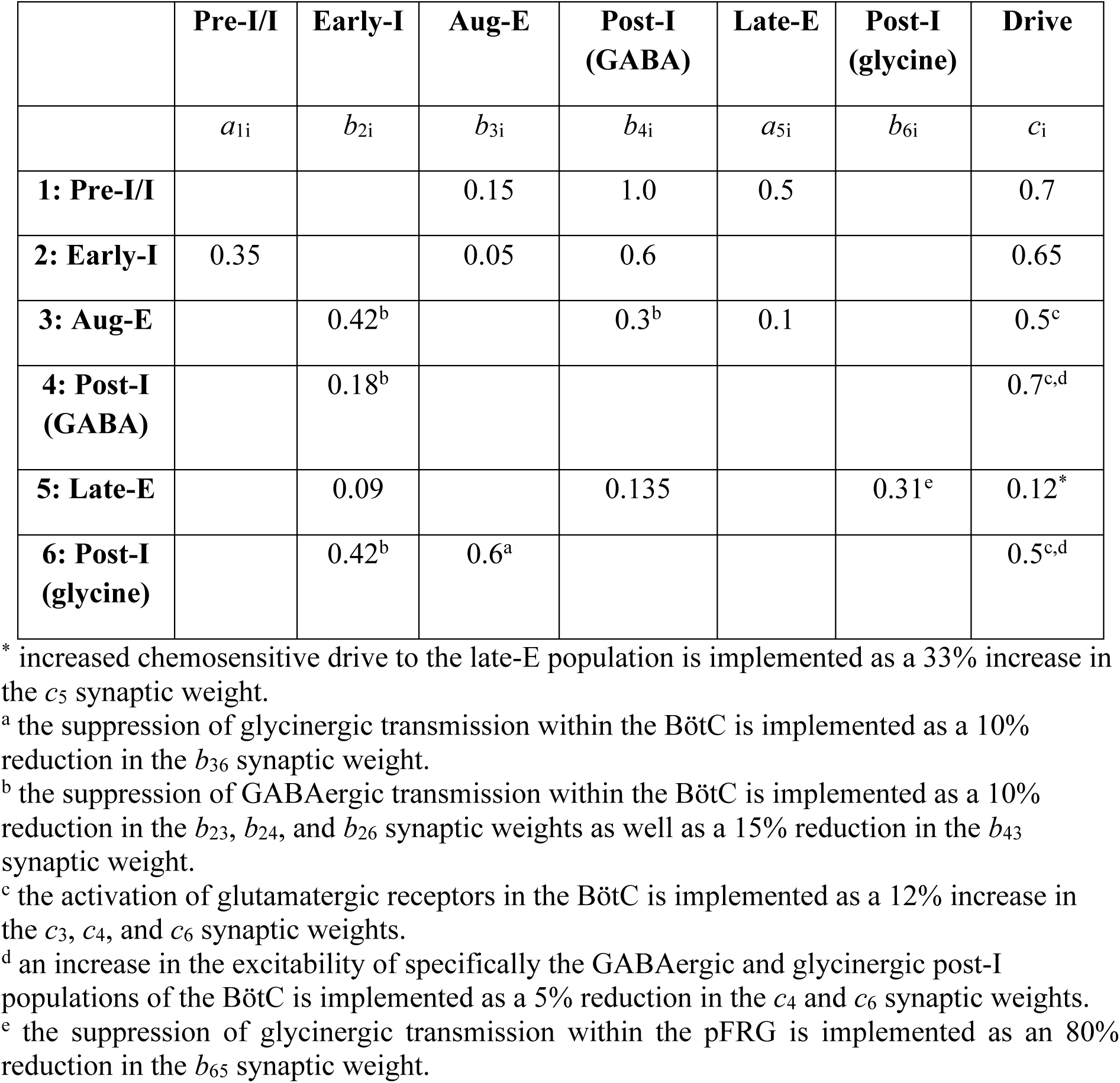
Summary of model connectivity and its manipulations performed to reproduce and predict experimental observations.

### Model description

We used a reduced conductance-based model to compute the average membrane potential *V*_*i*_, *i* ∈ {1, …, 6} (see Table 1) and firing rates of neuronal populations in the VRC closely following Rubin et al. (2009), Rubin et al. (2011). The current balance equation for each population was described by

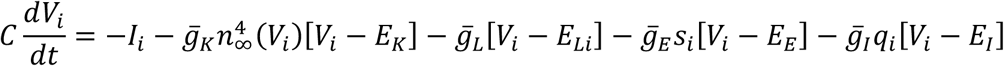

where 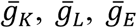, and 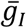 are the respective maximal conductances of a delayed rectifier potassium current, an ohmic leak current, an excitatory synaptic current, and an inhibitory synaptic current. The parameters *E*_*k*_, *E*_*Li*_, *E*_*E*_, and *E*_*I*_ are the reversal potentials of the potassium current, leak current, excitatory synaptic current, and inhibitory synaptic current. The voltage-dependent steady state activation of the potassium current is given by the expression *n*_∞_(*V*) = 1/[1 + exp (−[*V* + 30]/4)].

The current *I*_*i*_ differs by neuronal populations; it described either the persistent sodium current (*I*_*Nap*_) or an outward adapting current (*I*_*AD*_). The pre-I/I and late-E populations, (*i* = 1,5), contained 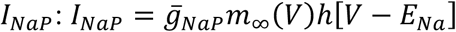. The parameters 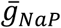 and *E*_*Na*_ were the maximal conductance of the persistent sodium current and the reversal potential of sodium. The activation of *I*_*Nap*_ was assumed instantaneous, and it was described by the steady state activation *m*_∞_(*V*) = 1/[1 + exp(−[*V* + 40]/6)]. The inactivation of *I*_*Nap*_ (*h*) was a dynamical variable governed by the following differential equation:

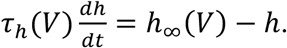

The steady state activation of *h* was described as *h*_∞_(*V*) = 1/[1 + exp([*V* + 55]/10)], and its time constant was specified by *τ*_*h*_(*V*) = 4000/cosh([*V* + 55]/10) ms.

The early-I, aug-E, and post-I_GABA_ populations, (*i* = 2,3,4), possessed an adaptive potassium current (*I*_*AD*_), which was described as 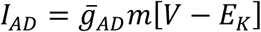. The parameter 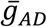 was the maximal conductance of the adaptive potassium current. Its activation (*m*) was a dynamical variable governed by the following differential equation:

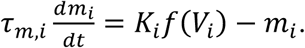

The steady state activation of *m* was determined by *K*_*i*_*f*(*V*_*i*_), where *K* was a scaling factor and *f*(*V*_*i*_) is the neuronal firing rate where *f*(*V*) was defined as

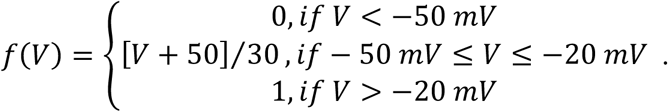

In the post-I_Gly_ population, (*i* = 6), there was no additional ionic current (*I*_6_ = 0).

The excitatory (*s*_*i*_) and inhibitory (*q*_*i*_) synaptic activations were determined by the activity of presynaptic populations as described by

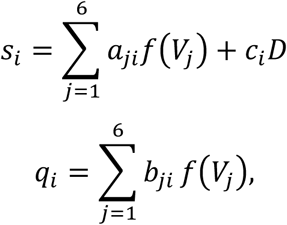

where *f*(*V*_*j*_) is the neuronal firing rate of the presynaptic population, and the synaptic weights *a, b*, and *c* can be found in Table 1 and depicted in Figure 5. The parameter *D* represented an external tonic drive, and it was equal to 1. Biophysical parameter values are described in Table 2.

**Table 2.**
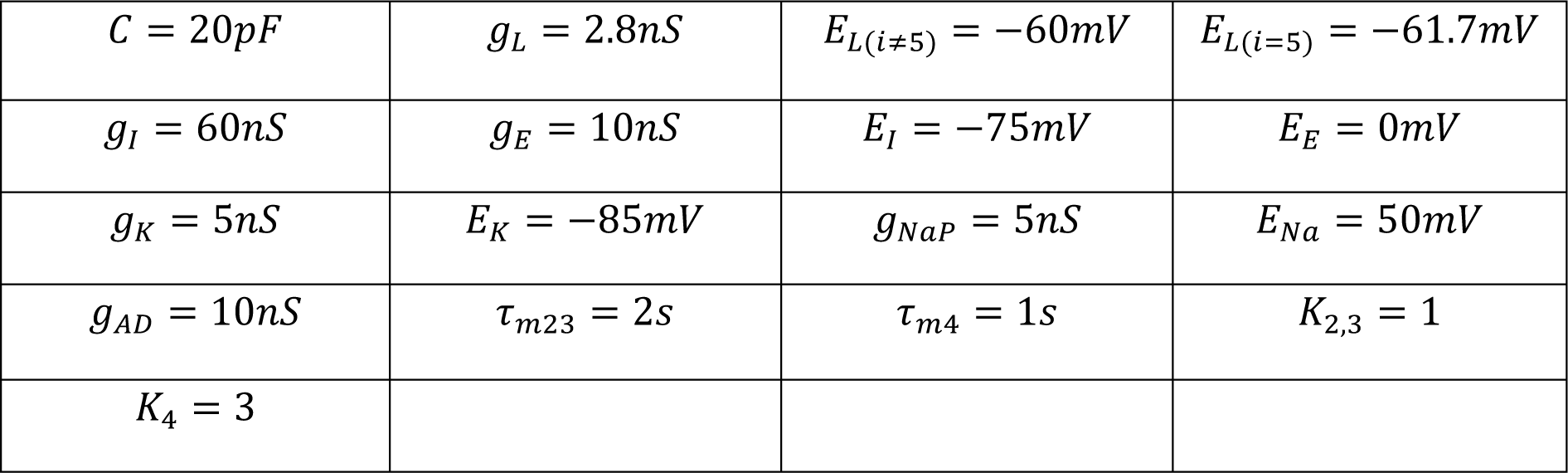
Biophysical parameters of the computational model of brainstem respiratory circuitry. The parameters 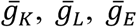 and 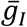 are the respective maximal conductances of a delayed rectifier potassium current, an ohmic leak current, an excitatory synaptic current, and an inhibitory synaptic current. The parameters *E*_*k*_, *E*_*Li*_, *E*_*E*_, and *E*_*I*_ are the reversal potentials of the potassium current, leak current, excitatory synaptic current, and inhibitory synaptic current. The parameters 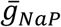 and *E*_*Na*_ are the maximal conductance of the persistent sodium current and the reversal potential of sodium. The parameters *τ*_*m,i*_ define the time constant of activation of an outward adapting current. The parameters *K*_*i*_ define a scaling factor in the activation on an outward adapting current.

**Table 3.** Average values of PN burst frequency (freq) and amplitude (amp), times of inspiration (Ti) and expirtion (Te), cVN post-I mean activity and duration, and mean activity of AbN during E1 and E2 phases of *in situ* preparations of control rats during baseline (normocapnia), hypercapnia (8% CO_2_), before and after bilateral microinjections of strychnine in the BötC (n=7).

|  | Baseline | Hypercapnia | Baseline<br>after strychnine | Hypercapnia<br>after strychnine |
| --- | --- | --- | --- | --- |
| PN freq (bpm) | 23±11 | 20±10* | 22±9 | 21±8 |
| PN amp (μV) | 30.6±10.2 | 31.6±14.8 | 22.8±10.3* | 23.6±10.1 |
| Ti (s) | 0.873±0.322 | 0.813±0.281* | 1.013±0.327 | 0.967±0.319<sup>#</sup> |
| Te (s) | 2.757±1.581 | 3.150±1.574* | 1.607±0.734* | 1.779±0.826 |
| cVN activity (μV) | 21.4±1.6 | 21.9±1.2 | 17.8±2.6* | 18.4±2.5 |
| cVN duration (% Te) | 75±10 | 71±8 | 60±17* | 57±14 |
| AbN during E1 (μV) | 3.2±1.5 | 3.0±1.6 | 2.6±1.6 | 2.4±1.0 |
| AbN during E2 (μV) | 6.4±1.5 | 9.4±1.3* | 4.7±2.1* | 6.0±2.6<sup>#†</sup> |

\* - different from baseline, # - different from baseline after strychnine, † - different from hypercapnia; P<0.05.

## Acknowledgements

This work was supported by São Paulo State Research Foundation (FAPESP, Grants # 2013/17.251-6 and 2015/23568-8), National Institutes of Health (Grants # R01AT008632 and U01EB021960) and Conselho Nacional de Desenvolvimento Científico e Tecnológico (CNPq, Grants # 310331/2017-0 and 408950/2018-8).

## Author contributions

KCF and WB, Conception and design, acquisition of data, analysis and interpretation of data, drafting or revising the article; MKA, acquisition of data, drafting or revising the article; YM and DBZ, Conception and design, analysis and interpretation of data, drafting or revising the article.

## Competing Interests statement

The authors declare no competing interests.

## REFERENCES

Abdala, A. P., Rybak, I. A., Smith, J. C. & Paton, J. F. 2009. Abdominal expiratory activity in the rat brainstem-spinal cord in situ: patterns, origins and implications for respiratory rhythm generation. J Physiol, 587, 3539–59.

Anderson, T. M., Garcia, A. J., 3rd, Baertsch, N. A., Pollak, J., Bloom, J. C., Wei, A. D., Rai, K. G. & Ramirez, J. M. 2016. A novel excitatory network for the control of breathing. Nature, 536, 76–80.

Anderson, T. M. & Ramirez, J. M. 2017. Respiratory rhythm generation: triple oscillator hypothesis. F1000Res, 6, 139.

Barnett, W. H., Abdala, A. P., Paton, J. F., Rybak, I. A., Zoccal, D. B. & Molkov, Y. I. 2017. Chemoreception and neuroplasticity in respiratory circuits. Exp Neurol, 287, 153–164.

Barnett, W. H., Jenkin, S. E. M., Milsom, W. K., Paton, J. F. R., Abdala, A. P., Molkov, Y. I. & Zoccal, D. B. 2018. The Kolliker-Fuse nucleus orchestrates the timing of expiratory abdominal nerve bursting. J Neurophysiol, 119, 401–412.

Bianchi, A. L., Denavit-Saubie, M. & Champagnat, J. 1995. Central control of breathing in mammals: neuronal circuitry, membrane properties, and neurotransmitters. Physiol Rev, 75, 1–45.

Bongianni, F., Mutolo, D., Cinelli, E. & Pantaleo, T. 2010. Respiratory responses induced by blockades of GABA and glycine receptors within the Botzinger complex and the pre-Botzinger complex of the rabbit. Brain Res, 1344, 134–47.

Bryant, T. H., Yoshida, S., De Castro, D. & Lipski, J. 1993. Expiratory neurons of the Botzinger Complex in the rat: a morphological study following intracellular labeling with biocytin. J Comp Neurol, 335, 267–82.

De Britto, A. A. & Moraes, D. J. 2017. Non-chemosensitive parafacial neurons simultaneously regulate active expiration and airway patency under hypercapnia in rats. J Physiol, 595, 2043–2064.

Del Negro, C. A., Funk, G. D. & Feldman, J. L. 2018. Breathing matters. Nat Rev Neurosci, 19, 351–367.

Dutschmann, M. & Herbert, H. 2006. The Kolliker-Fuse nucleus gates the postinspiratory phase of the respiratory cycle to control inspiratory off-switch and upper airway resistance in rat. Eur J Neurosci, 24, 1071–84.

Ezure, K. 1990. Synaptic connections between medullary respiratory neurons and considerations on the genesis of respiratory rhythm. Prog Neurobiol, 35, 429–50.

Ezure, K., Tanaka, I. & Kondo, M. 2003a. Glycine is used as a transmitter by decrementing expiratory neurons of the ventrolateral medulla in the rat. J Neurosci, 23, 8941–8.

Ezure, K., Tanaka, I. & Saito, Y. 2003b. Activity of brainstem respiratory neurones just before the expiration-inspiration transition in the rat. J Physiol, 547, 629–40.

Ezure, K., Tanaka, I. & Saito, Y. 2003c. Brainstem and spinal projections of augmenting expiratory neurons in the rat. Neurosci Res, 45, 41–51.

Flor, K. C., Silva, E. F., Menezes, M. F., Pedrino, G. R., Colombari, E. & Zoccal, D. B. 2018. Short-Term Sustained Hypoxia Elevates Basal and Hypoxia-Induced Ventilation but Not the Carotid Body Chemoreceptor Activity in Rats. Front Physiol, 9, 134.

Fortuna, M. G., Kugler, S. & Hulsmann, S. 2018. Probing the function of glycinergic neurons in the mouse respiratory network using optogenetics. Respir Physiol Neurobiol.

Geerling, J. C., Yokota, S., Rukhadze, I., Roe, D. & Chamberlin, N. L. 2017. Kolliker-Fuse GABAergic and glutamatergic neurons project to distinct targets. J Comp Neurol, 525, 1844–1860.

Guyenet, P. G. 2014. Regulation of breathing and autonomic outflows by chemoreceptors. Compr Physiol, 4, 1511–62.

Harris-Warrick, R. M. 2010. General principles of rhythmogenesis in central pattern generator networks. Prog Brain Res, 187, 213–22.

Huckstepp, R. T., Henderson, L. E., Cardoza, K. P. & Feldman, J. L. 2016. Interactions between respiratory oscillators in adult rats. Elife, 5.

Iceman, K. E. & Harris, M. B. 2014. A group of non-serotonergic cells is CO2-stimulated in the medullary raphe. Neuroscience, 259, 203–13.

Janczewski, W. A. & Feldman, J. L. 2006. Distinct rhythm generators for inspiration and expiration in the juvenile rat. J Physiol, 570, 407–20.

Jenkin, S. E. & Milsom, W. K. 2014. Expiration: breathing’s other face. Prog Brain Res, 212, 131–47.

Jenkin, S. E., Milsom, W. K. & Zoccal, D. B. 2017. The Kolliker-Fuse nucleus acts as a timekeeper for late-expiratory abdominal activity. Neuroscience, 348, 63–72.

Kam, K., Worrell, J. W., Janczewski, W. A., Cui, Y. & Feldman, J. L. 2013. Distinct inspiratory rhythm and pattern generating mechanisms in the preBotzinger complex. J Neurosci, 33, 9235–45.

Kubin, L., Alheid, G. F., Zuperku, E. J. & Mccrimmon, D. R. 2006. Central pathways of pulmonary and lower airway vagal afferents. J Appl Physiol, 101, 618–27.

Kumar Jha, P., Challet, E. & Kalsbeek, A. 2015. Circadian rhythms in glucose and lipid metabolism in nocturnal and diurnal mammals. Mol Cell Endocrinol, 418 Pt 1, 74–88.

Lemes, E. V., Aiko, S., Orbem, C. B., Formentin, C., Bassi, M., Colombari, E. & Zoccal, D. B. 2016. Long-term facilitation of expiratory and sympathetic activities following acute intermittent hypoxia in rats. Acta Physiol (Oxf), 217, 254–66.

Lemes, E. V. & Zoccal, D. B. 2014. Vagal afferent control of abdominal expiratory activity in response to hypoxia and hypercapnia in rats. Respir Physiol Neurobiol, 203, 90–7.

Lindsey, B. G., Rybak, I. A. & Smith, J. C. 2012. Computational models and emergent properties of respiratory neural networks. Compr Physiol, 2, 1619–70.

Marchenko, V., Koizumi, H., Mosher, B., Koshiya, N., Tariq, M. F., Bezdudnaya, T. G., Zhang, R., Molkov, Y. I., Rybak, I. A. & Smith, J. C. 2016. Perturbations of Respiratory Rhythm and Pattern by Disrupting Synaptic Inhibition within Pre-Botzinger and Botzinger Complexes. eNeuro, 3.

Marina, N., Abdala, A. P., Trapp, S., Li, A., Nattie, E. E., Hewinson, J., Smith, J. C., Paton, J. F. & Gourine, A. V. 2010. Essential role of Phox2b-expressing ventrolateral brainstem neurons in the chemosensory control of inspiration and expiration. J Neurosci, 30, 12466–73.

Molkov, Y. I., Abdala, A. P., Bacak, B. J., Smith, J. C., Paton, J. F. & Rybak, I. A. 2010. Late-expiratory activity: emergence and interactions with the respiratory CpG. J Neurophysiol, 104, 2713–29.

Molkov, Y. I., Rubin, J. E., Rybak, I. A. & Smith, J. C. 2016. Computational models of the neural control of breathing. Wiley Interdiscip Rev Syst Biol Med.

Molkov, Y. I., Shevtsova, N. A., Park, C., Ben-Tal, A., Smith, J. C., Rubin, J. E. & Rybak, I. A. 2014. A closed-loop model of the respiratory system: focus on hypercapnia and active expiration. PLoS One, 9, e109894.

Molkov, Y. I., Zoccal, D. B., Moraes, D. J., Paton, J. F., Machado, B. H. & Rybak, I. A. 2011. Intermittent hypoxia-induced sensitization of central chemoreceptors contributes to sympathetic nerve activity during late expiration in rats. Journal of neurophysiology, 105, 3080–91.

Moraes, D. J., Bonagamba, L. G., Costa, K. M., Costa-Silva, J. H., Zoccal, D. B. & Machado, B. H. 2014. Short-term sustained hypoxia induces changes in the coupling of sympathetic and respiratory activities in rats. J Physiol, 592, 2013–33.

Moraes, D. J., Bonagamba, L. G., Costa-Silva, J. H., Zoccal, D. B. & Machado, B. H. 2011. Glutamate receptors in ventral medulla are essential for expiratory and inspiratory responses to chemoreflex activation in unanesthetized rats. The FASEB Journal, 25, 1076.10.

Moraes, D. J., Dias, M. B., Cavalcanti-Kwiatkoski, R., Machado, B. H. & Zoccal, D. B. 2012a. Contribution of retrotrapezoid/parafacial respiratory region to the expiratory-sympathetic coupling in response to peripheral chemoreflex in rats. J Neurophysiol, 108, 882–890.

Moraes, D. J., Zoccal, D. B. & Machado, B. H. 2012b. Sympathoexcitation during chemoreflex active expiration is mediated by L-glutamate in the RVLM/Botzinger complex of rats. J Neurophysiol, 108, 610–23.

Pagliardini, S., Janczewski, W. A., Tan, W., Dickson, C. T., Deisseroth, K. & Feldman, J. L. 2011. Active expiration induced by excitation of ventral medulla in adult anesthetized rats. J Neurosci, 31, 2895–905.

Paton, J. F. 1996. A working heart-brainstem preparation of the mouse. J Neurosci Methods, 65, 63–8.

Paxinos, G. & Watson, C. 2007. The rat brain in stereotaxic coordinates, Amsterdam; Boston;, Academic Press/Elsevier.

Phillips, R. S., John, T. T., Koizumi, H., Molkov, Y. I. & Smith, J. C. 2019. Biophysical mechanisms in the mammalian respiratory oscillator re-examined with a new data-driven computational model. Elife, 8.

Powell, F. L., Milsom, W. K. & Mitchell, G. S. 1998. Time domains of the hypoxic ventilatory response. Respir Physiol, 112, 123–34.

Ramirez, J. M. & Baertsch, N. 2018. Defining the Rhythmogenic Elements of Mammalian Breathing. Physiology (Bethesda), 33, 302–316.

Ramirez, J. M., Telgkamp, P., Elsen, F. P., Quellmalz, U. J. & Richter, D. W. 1997. Respiratory rhythm generation in mammals: synaptic and membrane properties. Respir Physiol, 110, 71–85.

Richter, D. W. & Smith, J. C. 2014. Respiratory rhythm generation in vivo. Physiology, 29, 58–71.

Rubin, J. E., Bacak, B. J., Molkov, Y. I., Shevtsova, N. A., Smith, J. C. & Rybak, I. A. 2011. Interacting oscillations in neural control of breathing: modeling and qualitative analysis. J Comput Neurosci, 30, 607–32.

Rubin, J. E., Shevtsova, N. A., Ermentrout, G. B., Smith, J. C. & Rybak, I. A. 2009. Multiple Rhythmic States in a Model of the Respiratory Central Pattern Generator. Journal of Neurophysiology, 101, 2146–2165.

Rybak, I. A., Molkov, Y. I., Jasinski, P. E., Shevtsova, N. A. & Smith, J. C. 2014. Rhythmic bursting in the pre-Botzinger complex: mechanisms and models. Prog Brain Res, 209, 1–23.

Saper, C. B. 2006. Staying awake for dinner: hypothalamic integration of sleep, feeding, and circadian rhythms. Prog Brain Res, 153, 243–52.

Shevtsova, N. A. & Rybak, I. A. 2016. Organization of flexor-extensor interactions in the mammalian spinal cord: insights from computational modelling. J Physiol, 594, 6117–6131.

Smith, J. C., Abdala, A. P., Koizumi, H., Rybak, I. A. & Paton, J. F. 2007. Spatial and functional architecture of the mammalian brain stem respiratory network: a hierarchy of three oscillatory mechanisms. J Neurophysiol, 98, 3370–87.

Smith, J. C., Ellenberger, H. H., Ballanyi, K., Richter, D. W. & Feldman, J. L. 1991. Pre-Botzinger complex: a brainstem region that may generate respiratory rhythm in mammals. Science, 254, 726–9.

St-John, W. M., Stornetta, R. L., Guyenet, P. G. & Paton, J. F. 2009. Location and properties of respiratory neurones with putative intrinsic bursting properties in the rat in situ. J Physiol, 587, 3175–88.

Tan, W., Janczewski, W. A., Yang, P., Shao, X. M., Callaway, E. M. & Feldman, J. L. 2008. Silencing preBotzinger complex somatostatin-expressing neurons induces persistent apnea in awake rat. Nat Neurosci, 11, 538–40.

Tanaka, I., Ezure, K. & Kondo, M. 2003. Distribution of glycine transporter 2 mRNA-containing neurons in relation to glutamic acid decarboxylase mRNA-containing neurons in rat medulla. Neurosci Res, 47, 139–51.

Tian, G. F., Peever, J. H. & Duffin, J. 1999. Botzinger-complex, bulbospinal expiratory neurones monosynaptically inhibit ventral-group respiratory neurones in the decerebrate rat. Exp Brain Res, 124, 173–80.

Yang, C. F. & Feldman, J. L. 2018. Efferent projections of excitatory and inhibitory preBotzinger Complex neurons. J Comp Neurol.

Zoccal, D. B., Silva, J. N., Barnett, W. H., Lemes, E. V., Falquetto, B., Colombari, E., Molkov, Y. I., Moreira, T. S. & Takakura, A. C. 2018. Interaction between the retrotrapezoid nucleus and the parafacial respiratory group to regulate active expiration and sympathetic activity in rats. Am J Physiol Lung Cell Mol Physiol.

Zoccal, D. B., Simms, A. E., Bonagamba, L. G., Braga, V. A., Pickering, A. E., Paton, J. F. & Machado, B. H. 2008. Increased sympathetic outflow in juvenile rats submitted to chronic intermittent hypoxia correlates with enhanced expiratory activity. J Physiol, 586, 3253–65.

